# Noncanonical CD11b on CD8^+^ T cells promotes cytotoxic effector function

**DOI:** 10.64898/2026.10.07.757268

**Authors:** Jone Etxaniz-Diaz de Durana, Alexandre Bosch, Endika Prieto-Fernández, Lorena Pérez-Gutiérrez, Kexin Yu, Paloma Velasco-Beltrán, Leire Egia-Mendikute, Iker Oyenarte, So Young Lee, Ander de Blas, Iñaki Eguren-Santamaría, Jesús Jiménez-Barbero, Miguel F Sanmamed, Maria M Caffarel, Ana M. Aransay, Alena Gros, Pablo F. Cespedes, Asis Palazon, June Ereño-Orbea, Asier Antoñana-Vildosola

## Abstract

Cancer immunosurveillance largely relies on effector CD8^+^ T cells, yet surface determinants of cytotoxic competence within tumor-reactive populations remain incompletely defined. Using single-cell surface proteomics in a mouse breast cancer model, we identified a tumor-enriched subset of antigen-experienced CD8^+^ T cells expressing CD11b, a prototypic myeloid integrin. CD11b⁺ CD8⁺ T cells were detected across mouse tumor models and human solid tumors and, within tumor-reactive populations, CD11b marked cells with greater cytotoxic potential. Following activation, naive CD8^+^ T cells induced *ITGAM* transcription and CD18-dependent CD11b surface expression. Mechanistically, CD11b localized to the peripheral adhesive ring of the immune synapse and promoted lytic-granule secretion. Accordingly, CD11b^+^ T cells exhibited enhanced TCR- and CAR-mediated tumor cell killing, whereas CD11b deletion impaired T cell antitumor activity. Together, these findings identify CD11b as an inducible, noncanonical component of the T cell immune synapse that marks cytotoxic competence and promotes antitumor T cell function.

## INTRODUCTION

Cytotoxic CD8⁺ T lymphocytes (CTLs) are central effectors of antitumor immunity, yet tumor-infiltrating lymphocytes (TILs) are highly heterogeneous in both antigen specificity and functional state. Solid tumors contain tumor-reactive CD8⁺ T cells alongside abundant bystander populations, while tumor-reactive cells themselves encompass distinct differentiation and effector states^1, 2^. Co-expression of surface markers such as PD-1, CD39, and CD103 can enrich for tumor-reactive TILs^3, 4^. However, these phenotypically defined compartments remain functionally heterogeneous and do not fully resolve cytotoxic competence^5^. Identifying surface features that distinguish effector capacity within tumor-reactive CD8⁺ T cells is therefore important both for understanding endogenous tumor control and for selecting potent cells for adoptive T cell therapy. CTLs mediate target cell killing through polarized secretion of lytic granules at the cytotoxic immune synapse^6^. Following T cell receptor (TCR) recognition of peptide–MHC (pMHC) class I complexes, this interface organizes into supramolecular activation clusters that coordinate signaling, adhesion and cytotoxic secretion^7, 8^. The β2 integrin LFA-1 (CD11a–CD18; αLβ2) is central to this architecture, stabilizing T cell–target cell interactions and directing lytic-granule release^9, 10^. CD18 can also pair with CD11b (αMβ2; Mac-1), CD11c (αXβ2), or CD11d (αDβ2) to form three alternative β2 integrins, but their roles in T cell cytotoxicity remain poorly understood. Recent evidence that unconventional integrin pairing can potentiate TCR signaling and antigen-specific cytotoxicity suggests that the integrin repertoire of activated T cells is more dynamic than previously appreciated^11^. Whether T cells deploy alternative β2 integrins at the cytotoxic immune synapse remains unknown.

Here, using high-dimensional cell-surface protein profiling, we identify a tumor-enriched subset of antigen-experienced CD8⁺ T cells with non-canonical expression of integrin CD11b. CD11b^+^ CD8^+^ T cells were present across multiple syngeneic murine tumor models and human solid tumors, and CD11b expression marked enhanced effector capacity within tumor-reactive TIL and CAR-T cell populations. TCR activation induced CD11b expression in CD8^+^ T cells, and CD11b localized to the cytotoxic immune synapse, where it promoted lytic-granule release and antigen-specific tumor-cell killing. Together, our findings establish activation-induced remodeling of the β2-integrin repertoire as a regulator of cytotoxic T cell function.

## RESULTS

### Surface protein profiling identifies CD11b^+^ CD8^+^ TILs in solid tumors

To define surface-receptor heterogeneity during antitumor immunity, we performed high-dimensional flow cytometry profiling on TILs isolated from orthotopic EO771 breast tumors. TILs were analyzed alongside splenic lymphocytes from tumor-bearing mice, as well as mammary fat pad (MFP) and spleen tissues from tumor-free controls. Samples were tissue-barcoded and stained with a common backbone panel to enable consistent identification of immune populations across tissues (**Fig. 1a, Extended Data Fig. 1a, and Methods**). To identify surface markers associated with antitumor CD8^+^ T cell responses, we focused on antigen-experienced (CD44^+^) CD8^+^ T cells (**Extended Data Fig. 1b**). Of the 255 surface markers evaluated, 121 were detected on ≥10% of cells in at least one tissue and retained for analysis. Among these, we identified 49 that were differentially expressed (fold-change >2), of which 45 were increased in tumor-infiltrating CD8^+^ T cells and 4 in non-tumor tissues **(Fig. 1b–c).** Antigen-experienced CD8^+^ TILs showed a substantial increase in CD11b (integrin αM), a β2 integrin classically associated with myeloid cells^12^. This striking enrichment of CD11b on tumor-infiltrating CD8^+^ TILs prompted us to investigate CD11b⁺ CD8⁺ T cells further.

**Figure 1.**
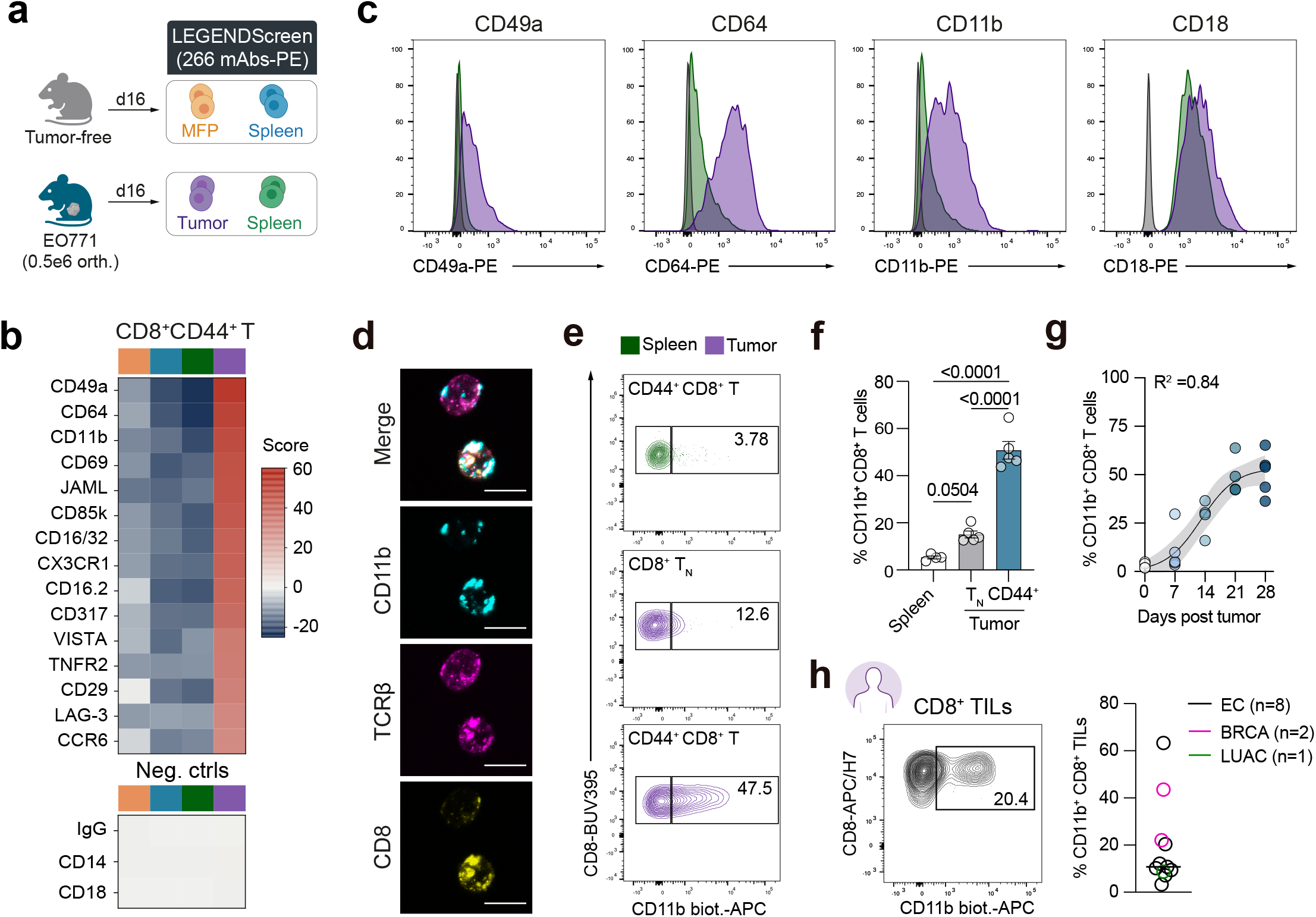
Surface protein profiling identifies CD11b^+^ CD8^+^ TILs in solid tumors. **a,** Schematic overview of the CD8^+^ T cell surface marker screen. EO771 tumors and spleens from tumor-bearing mice and mammary fat pads (MFPs) and spleens from tumor-free mice were collected 16 d after tumor implantation (n = 4 per group, except n = 10 tumor-free MFPs). Samples were pooled within tissue group, barcoded, stained with a common backbone antibody panel, and distributed across the LEGENDScreen Mouse PE panel. **b,** Top 15 surface markers and negative controls (Neg. ctrls) expressed by antigen-experienced CD44⁺CD8⁺TCRβ⁺ T cells from EO771 tumors relative to the three non-tumor tissues. Colors indicate tumor-free MFP (orange), tumor-free spleen (blue), spleen from tumor-bearing mice (green), and EO771 tumor (purple). **c,** Representative histograms of selected markers in EO771 tumor-infiltrating (purple) and splenic (green) antigen-experienced CD8⁺ T cells, and corresponding isotype controls (gray). **d,** Representative confocal image of FACS-purified CD11b⁺CD8⁺ TILs from EO771 tumors stained for CD11b (cyan), TCRβ (magenta), and CD8 (yellow). Scale bar, 10 μm. **e,f,** Representative contour plots (**e**) and frequency (**f**) of CD11b^+^ cells among antigen-experienced (CD44^+^) splenic CD8⁺ T cells, intratumoral naive CD8^+^ T cells (CD44⁻CD62L⁺), and antigen-experienced CD8^+^ TILs from EO771 tumor-bearing mice (n = 4–5 mice). TN, naive CD8^+^ T cells; Spl., spleen. **g,** Frequency of CD11b⁺ cells among antigen-experienced CD8⁺ TILs at the indicated time points during EO771 tumor progression (n = 3–5 mice per time point). Temporal changes were fitted by nonlinear regression using a logistic growth model; the solid line indicates the best-fit curve and shading indicates the 95% confidence interval. **h,** Representative flow cytometry plot (left) and quantification (right) of CD11b expression among CD8⁺ TILs from patients with endometrial cancer (EC; n = 8 patients, black), lung adenocarcinoma (LUAC; n = 1 patient, green), and breast cancer tumors (BRCA; n = 2 patients, magenta). *P* values were calculated using one-way analysis of variance (ANOVA) with Tukey’s multiple-comparison test (**f**).

We first confirmed CD11b expression on tumor-infiltrating CD8^+^ T cells. Confocal imaging of FACS-purified CD11b^+^ CD8^+^ TILs confirmed co-expression of CD11b, TCRβ, and CD8 on individual cells **(Fig. 1d)**. Consistent with the predominant expression of CD11b in the myeloid cell compartment, CD11b surface abundance on CD8⁺ TILs was approximately tenfold lower than on tumor-infiltrating myeloid cells. CD11b was also detected on subsets of tumor-infiltrating CD4⁺ T cells, B cells, and NK cells **(Extended Data Fig. 1c–d)**, suggesting that CD11b expression extends to several lymphoid populations in the tumor microenvironment.

We next examined CD11b expression across subsets of CD8^+^ T cells in EO771 tumor-bearing mice. Consistent with our initial screen, a substantial fraction of antigen-experienced CD8^+^ TILs expressed CD11b **(Fig. 1e-f, Extended Data Fig. 2a)**. By contrast, CD11b expression was largely absent from intratumoral naive (CD62L^+^CD44^−^) CD8^+^ T cells (CD8^+^ TN) and splenic antigen-experienced CD8^+^ T cells from tumor-bearing mice **(Fig. 1e–f)**. Longitudinal analysis of EO771 tumors showed that antigen-experienced CD11b^+^ CD8^+^ TILs emerged within one week of tumor implantation and increased in frequency during tumor progression (**Fig. 1g, Extended Data Fig. 2b).**

We next assessed whether CD11b expression on CD8^+^ TILs was conserved across tumor models. CD11b^+^ CD8^+^ TILs were detected in YUMMER1.7 and B16-OVA melanoma tumors and in the ID8-OVA ovarian cancer model **(Extended Data Fig. 2c–e)**. Moreover, CD11b was detected on a fraction of CD8^+^ TILs from patients with endometrial, breast, and lung tumors **(Fig. 1h and Extended Data Fig. 2f)**. Thus, surface expression of CD11b identifies a subset of antigen-experienced CD8^+^ TILs across multiple mouse tumor models and human solid tumors.

### CD11b identifies highly functional tumor-reactive CD8^+^ T cells

Because PD-1 is an established surrogate marker of tumor-reactive CD8^+^ T cells^3^, we compared PD-1 expression between CD11b^+^ and CD11b^−^ CD8^+^ TILs from EO771 tumors. Notably, CD11b^+^ CD8^+^ TILs contained a significantly higher frequency of PD-1^+^ cells than their CD11b^−^ counterparts (**Fig. 2a**), suggesting that CD11b is preferentially expressed within CD8^+^ T cell populations enriched for tumor reactivity.

**Figure 2.**
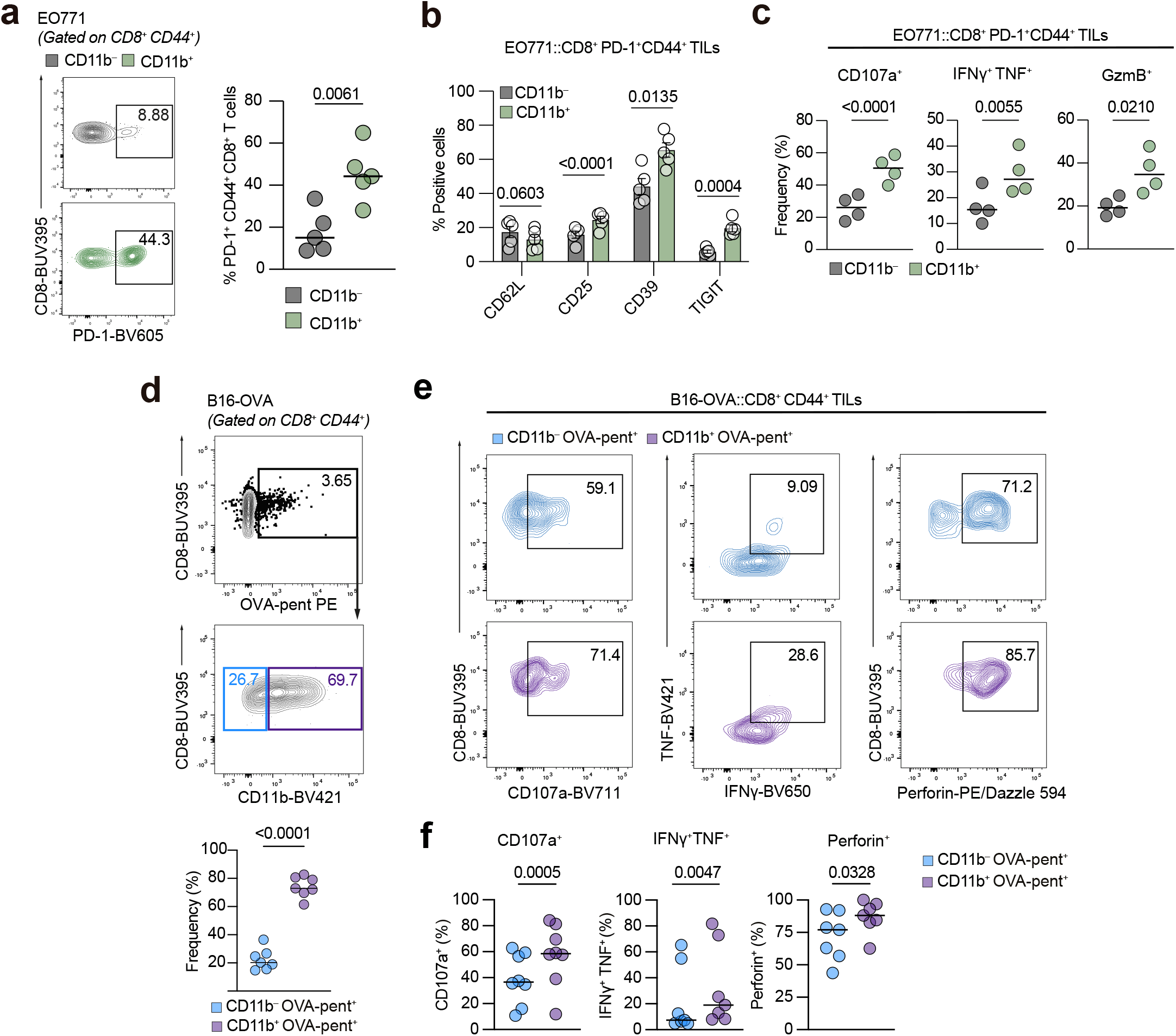
CD11b identifies an effector-enriched state within tumor-reactive CD8⁺ T cells. **a,** Representative contour plots and quantification of PD-1⁺ cells among CD11b⁻ (top) or CD11b⁺ (bottom) antigen-experienced CD8⁺ T cells from EO771 tumors (n = 5 mice). **b,** Frequency of cells expressing the indicated markers within CD11b⁻PD-1⁺ or CD11b⁺PD-1⁺ CD8⁺ TILs from EO771 tumors (n = 5 mice). **c,** Frequencies of CD107a^+^, IFN-γ^+^TNF^+^ and GzmB⁺ cells within CD11b⁻PD-1⁺ and CD11b⁺PD-1⁺ CD8⁺ TILs from EO771 tumors after overnight ex vivo stimulation (n = 4 mice). **d,** Representative contour plots and quantification of CD11b expression among OVA-pentamer^+^ CD8^+^ TILs from B16-OVA tumors collected on day 21 (n = 6 mice). **e**, **f**, Representative plots (**e**) and quantification (**f**) of CD107a^+^, IFN-γ^+^TNF^+^ and perforin^+^ cells within CD11b⁻ or CD11b⁺ OVA-pentamer⁺ CD8⁺ TILs from B16-OVA tumors after overnight ex vivo stimulation (n = 6 mice). Data are represented as mean ± s.e.m. (**b**). P values were calculated using two-tailed, paired Student’s t test (**a**–**d, f**).

Given the known functional heterogeneity of PD-1^+^ TILs^5^, we next asked whether CD11b further stratified this population. Within PD-1^+^ CD8^+^ TILs, CD11b^+^ cells exhibited a distinct phenotype, with higher levels of activation markers such as CD25, CD39, and TIGIT, compared to CD11b^−^PD-1^+^ CD8^+^ T cells (**Fig. 2b**). Functionally, the CD11b^+^ PD-1^+^ CD8^+^ TIL subset exhibited higher frequencies of cells undergoing degranulation (CD107a) and expressing IFN-γ, TNF, and granzyme B (GzmB) following ex vivo stimulation **(Fig. 2c, Extended Data Fig. 3a)**. These findings suggest that CD11b may provide a phenotypic strategy to enrich for highly functional CD8^+^ TILs with tumor-reactive features.

To directly assess CD11b expression within tumor-reactive CD8^+^ T cells, we engineered B16, EO771, and ID8 cancer cell lines to express the model antigen ovalbumin (OVA) and detected OVA257-264 (SIINFEKL)-specific CD8⁺ T cells using H2-Kᵇ-SIINFEKL pentamers. Tumor-infiltrating and circulating pentamer^+^ CD8⁺ T cells included a fraction of CD11b^+^ CD8^+^ T cells across all three models (**Fig. 2d, Extended Data Fig. 3b–c).** Within OVA-specific CD8⁺ TILs from B16-OVA tumors, CD11b⁺ cells expressed higher levels of multiple activation markers compared to their CD11b⁻ counterparts **(Extended Data Fig. 3d).** Beyond phenotypic differences, we also observed distinct functional capacities between those two tumor-reactive CD8^+^ T cell subsets. Upon ex vivo restimulation, the OVA-pentamer^+^ CD11b^+^ CD8^+^ T cells contained higher frequencies of cells undergoing degranulation (CD107a) and expressing IFN-γ, TNF, and perforin compared to their CD11b⁻ counterparts **(Fig. 2e–f)**. These findings identify CD11b as a candidate surface marker of cytotoxic competence within antigen-specific CD8⁺ T cell populations.

### CD11b is induced upon T cell activation and recruited to the immune synapse

Because CD11b is abundant on myeloid cells and T cells can acquire membrane proteins from interacting cells^13, 14, 15^, we first asked whether CD8^+^ T cells intrinsically express CD11b. Purified naive human CD8^+^ T cells had undetectable *ITGAM* (encoding CD11b) or *ITGAX* (encoding CD11c) transcripts, but constitutively expressed *ITGAL* (encoding CD11a) and *ITGB2* (encoding CD18) mRNA. Following polyclonal activation, *ITGAM* and *ITGB2* transcript abundance increased significantly, whereas *ITGAL* and *ITGAX* mRNA remained unchanged **(Fig. 3a, Extended Data Fig. 4a)**. Consistent with this transcriptional induction, CD11b surface expression became detectable after 96 h of T cell activation and increased progressively with subsequent cell division rounds in activated human **(Fig. 3b–c)** and mouse **(Extended Data Fig. 4b)** CD8^+^ T cells. Over the same activation time course, CD3 and CD11a underwent an early decrease in surface abundance followed by recovery^16, 17^ while CD11b progressively accumulated over the same time period **(Fig. 3b,c)**. Thus, T cell activation remodels the β2-integrin repertoire of CD8⁺ T cells through de novo induction of CD11b.

**Figure 3.**
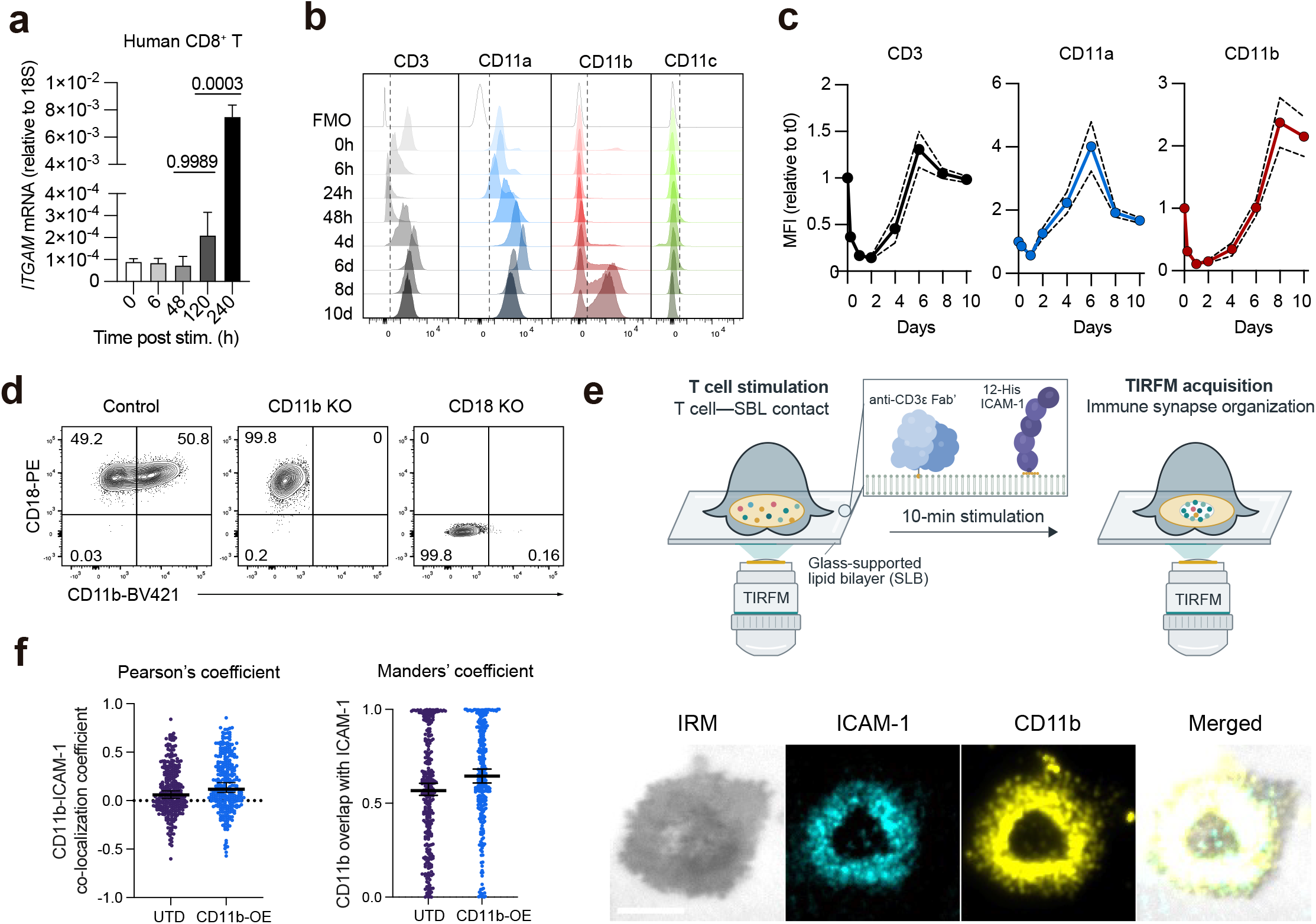
CD11b is induced after T cell activation and recruited to the immune synapse. **a**, *ITGAM* transcript abundance measured by qPCR in naive human CD8⁺ T cells at the indicated times after in vitro activation (n = 3 donors). **b**, **c,** Representative flow cytometry histograms of CD3, CD11a, CD11b, and CD11c surface expression during a 10-day activation time course (**b**) and gMFI of CD3, CD11a, and CD11b relative to day 0 (**c**) during the same activation time course (n = 3 donors). **d,** Representative flow cytometry plots showing CD11b surface expression in control, CD11b-deficient and CD18-deficient human CD8⁺ T cells. Representative of three independent experiments. **e,** Schematic of the glass-supported lipid bilayers (SLB) assay used to visualize CD11b distribution at the T cell immune synapse (top) and representative total internal reflection fluorescence microscopy (TIRFM) images acquired after 10 min of stimulation (bottom). CD8^+^ T cells were stimulated on SLBs functionalized with monomeric anti-CD3ε Fab’ and ICAM-1. Images show the internal reflection microscopy (IRM) footprint, ICAM-1 (cyan), CD11b (yellow), and merged channels. Scale bar = 5 µm. **f**, Pearson correlation coefficient and Manders’ colocalization coefficient quantifying, respectively, pixel-intensity correlation between CD11b and ICAM-1 and the fraction of CD11b signal overlapping ICAM-1 within individual T cell synapses formed by UTD or CD11b-overexpressing (CD11b OE) T cells. Each point represents one synapse; horizontal lines indicate medians. P values were calculated using one-way analysis of variance (ANOVA) with Tukey’s multiple-comparison test (**a**).

Surface expression of integrin α chains requires association with a β subunit. We therefore asked whether activation-induced CD11b on CD8^+^ T cells formed the canonical αMβ2 heterodimer with CD18. Genetic knockout of CD18 using CRISPR/Cas9 abolished CD11b surface expression in activated human CD8^+^ cells **(Fig. 3d)**, demonstrating that CD11b surface expression depends on CD18. CD11a also pairs with CD18 and is expressed at substantially higher levels in T cells, raising the possibility that competition for CD18 might constrain CD11b surface availability. However, CD11a knockout did not alter CD11b surface expression despite reducing CD18 levels **(Extended Data Fig. 4c)**, indicating that CD11b membrane expression is not limited by LFA-1 pairing.

Having established that activation-induced CD11b forms the canonical αMβ2 heterodimer, we next examined whether CD11b expressed on CD8^+^ T cells retains the cation-sensitive conformational regulation characteristic of β2 integrins^18^. Under resting conditions, CD11b predominantly adopted an inactive conformation, whereas Mn²⁺ promoted an active αI-domain state, as indicated by exposure of the activation-dependent CBRM1/5 epitope. Consistent with this, the I332G substitution, which favors the high-affinity conformation of CD11b^19, 20^, similarly stabilized CD11b active conformation in the absence of Mn²⁺ **(Extended Data Fig. 4d).** Thus, CD11b expressed on CD8^+^ T cells shows an activation-dependent conformational switching characteristic of functional β2 integrins.

Given the central role of β integrin organization in T cell adhesion and immune synapse function^10, 11, 21^, we next asked whether activation-induced CD11b is recruited to the immune synapse. To that end, in vitro expanded human T cells were stimulated on supported lipid bilayers (SLBs) functionalized with anti-CD3ε Fab and ICAM-1 and imaged using total internal reflection fluorescence microscopy (TIRFM). After 10 min of stimulation, endogenous CD11b on untransduced (UTD) T cells accumulated in ICAM-1-enriched peripheral regions of the synapse, consistent with localization to the peripheral supramolecular activation cluster (pSMAC) **(Fig. 3e–f and Extended Data Fig. 4e).** Similar pSMAC enrichment was observed in T cells overexpressing wild-type CD11b (**Fig. 3f and Extended Data Fig. 4e)**, which was independently confirmed using GFP-tagged CD11b **(Extended Data Fig. 4f).** Together, these findings establish activation-induced CD11b as a component of the peripheral adhesive ring of the CD8⁺ T cell immune synapse.

### CD11b promotes lytic granule secretion at the T cell immune synapse

To determine whether CD11b expression was associated with a distinct transcriptional state following T cell activation, we performed RNA-sequencing (RNA-seq) on sorted CD11b⁺ and CD11b⁻ human CD8⁺ T cells 24 h after polyclonal TCR restimulation. We identified 928 differentially expressed genes between the two populations **(Fig. 4a)**. CD11b⁺ CD8⁺ T cells expressed numerous genes associated with cytotoxic effector function, including *CX3CR1*, *KLRG1*, *PRF1*, and *IFNG*. By contrast, CD11b⁻ cells displayed increased expression of genes associated with stem- and memory-like T cell states, including *TCF7*, *LEF1*, and *IL7R* **(Fig. 4b).** Gene ontology (GO) Biological Process over-representation analysis and gene set enrichment analyses (GSEA) further showed that CD11b^+^ CD8^+^ T cells were enriched for immune effector programs, including cell killing and exocytosis, together with a human CD8 cytotoxic signature **(Fig. 4c–d).** Together with the reported role of CD11b in target-cell killing in other leukocyte populations^22, 23^, these data prompted us to ask whether CD11b regulates lytic granule secretion by T cells. To test this, we used TIRFM to quantify synaptic release of the lytic granule components perforin and granzyme B (GzmB) in T cell–SLB contacts formed by UTD control or CD11b-overexpressing T cells. CD11b overexpression significantly increased synaptic release of both perforin and GzmB relative to control T cells **(Fig. 4e–f),** demonstrating that CD11b expression is sufficient to enhance lytic granule secretion by T cells at the immune synapse.

**Figure 4.**
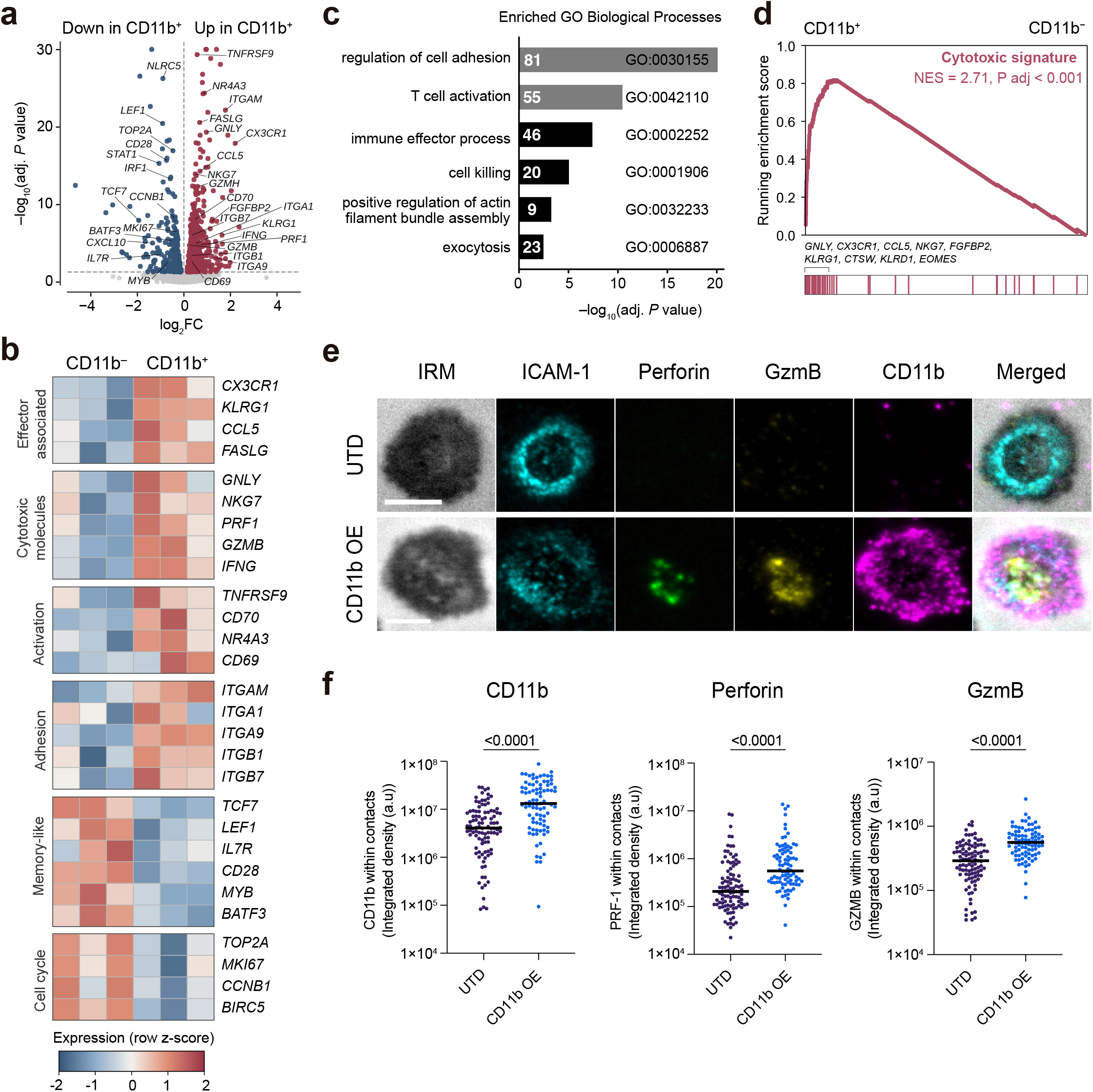
CD11b expression is associated with a cytotoxic T cell effector program and promotes lytic granule secretion. **a,** Volcano plot of differential gene expression between paired CD11b⁺ and CD11b⁻ human CD8⁺ T cells 24 h after TCR restimulation (n = 3 donors), showing log2 fold change versus −log10(adjusted *P* value). Differentially expressed genes (Benjamini–Hochberg-adjusted *P* < 0.05; red, higher expression in CD11b⁺ CD8⁺ T cells; blue, higher expression in CD11b⁻ CD8⁺ T cells) are indicated. Adjusted P values were capped for visualization. **b,** Heatmap of selected differentially expressed genes grouped by the indicated functional categories. Color scale represents row-normalized *z*-scores. **c,** GO Biological Process enrichment analysis of genes upregulated in CD11b⁺ cells, showing selected non-redundant terms. Terms related to cytotoxic effector function and granule exocytosis are shown in black; numbers indicate gene counts. **d,** GSEA of the published human CD8⁺ T-cell Cytotoxic signature^44^ in CD11b⁺ versus CD11b⁻ cells. Selected leading-edge genes are indicated. NES, normalized enrichment score. **e,** Representative TIRFM and internal reflection microscopy (IRM) images of synapses formed by untransduced (UTD) and CD11b-overexpressing (CD11b OE) human T cells on supported lipid bilayers presenting anti-CD3ε Fab′ and ICAM-1. Channels show the IRM footprint, ICAM-1 (cyan), secreted perforin (green) and GzmB (yellow), CD11b (magenta), and merged channels. Scale bars = 5 μm. **f,** Quantification of CD11b, GzmB and perforin integrated density within individual synapses formed by UTD and CD11b OE T cells. Each point represents one synapse sampled across three independent experiments; horizontal lines indicate medians. P values were calculated using a two-tailed Mann–Whitney U test.

### CD11b expression enhances T cell cytotoxicity

Having established that CD11b marks a highly functional tumor-reactive CD8⁺ T cell subset and regulates lytic granule secretion, we next sought to examine whether CD11b regulates antigen-specific CD8^+^ T cell effector function. We first compared the antigen-dependent killing capacity of CD11b⁺ and CD11b^−^ OVA-specific OT-I CD8⁺ T cells against B16-OVA target cells using real-time cytotoxicity assays. CD11b⁺ OT-I T cells mediated faster and more efficient target cell killing compared with their CD11b^−^ OT-I counterparts **(Fig. 5a–b and Extended Data Fig. 5a)**, suggesting that CD11b marks a T cell state poised for strong antigen-specific cytotoxic function.

**Figure 5.**
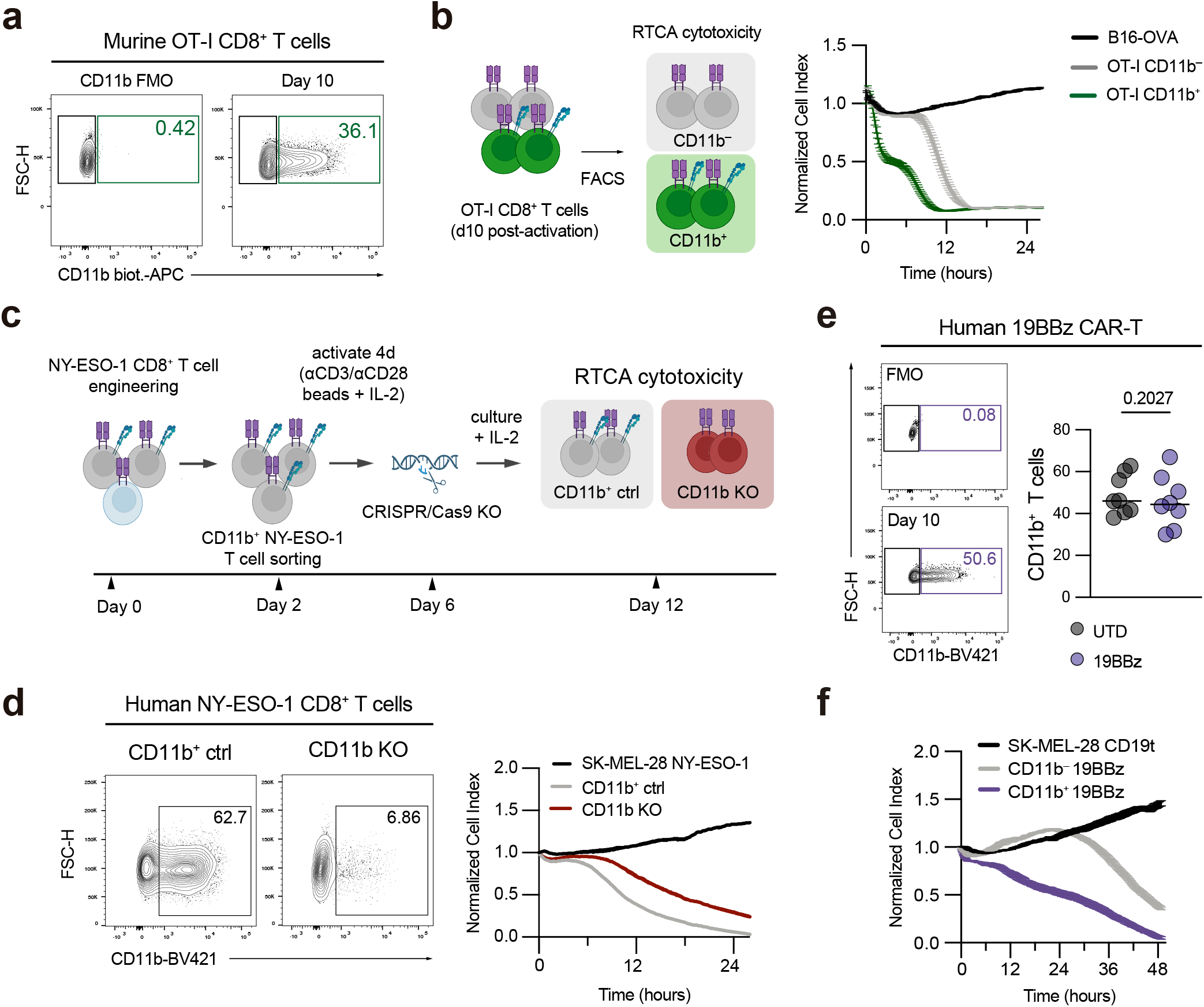
CD11b enhances antigen-specific T cell cytotoxicity. **a,** Representative flow cytometry plot of CD11b expression on OT-I CD8^+^ T cells 10 days after in vitro stimulation. Representative of four independent experiments. **b,** Experimental schematic (left) and real-time cytotoxicity assay (xCELLigence; right) evaluating B16-OVA tumor cell lysis when cultured alone or treated OT-I CD11b⁻ or OT-I CD11b⁺ CD8^+^ T cells (E:T = 0.25:1). Representative of three independent experiments; replicate-level data are shown in Extended Data Fig. 5a. **c,** Experimental workflow for generating CD11b-deficient NY-ESO-1-specific human CD8⁺ T cells. Cells were transduced with an NY-ESO-1-specific TCR, selected for CD11b expression, edited using CRISPR–Cas9, expanded and assessed for antitumor activity in real-time cytotoxicity assays. **d,** Representative flow cytometry plots showing CD11b expression in control-electroporated (CD11b^+^ ctrl) and ITGAM-targeted (CD11b KO) NY-ESO-1-specific CD8^+^ T cells (left), and real-time cytotoxicity assay (xCELLigence; right) evaluating SK-MEL28-NY-ESO-1 tumor cell lysis when cultured alone, or with CD11b^+^ ctrl or CD11b KO NY-ESO-1 CD8^+^ T cells (E:T = 0.25:1). Representative of three independent experiments; CD11b knockout efficiency and two additional independent cytotoxicity experiments are shown in Extended Data Fig. 5b and c, respectively. **e,** Representative flow cytometry plot (left) and quantification (right) of CD11b expression in untransduced (UTD) or 19BBz cells after 10 days of in vitro expansion (n = 7 donors). **f,** Real-time cytotoxicity assay (xCELLigence) evaluating SK-MEL28-CD19t tumor cell lysis when cultured alone or with CD11b^+^ or CD11b⁻ 19BBz CAR-T cells (E:T = 0.25:1). Representative of three independent experiments; replicate-level data are shown in Extended Data Fig. 5f. P values were calculated using two-tailed, paired Student’s t test (**e**).

To test whether CD11b directly contributes to cytotoxic T cell function, we followed a CRISPR/Cas9 approach to generate CD11b-knockout human CD8⁺ T cells expressing a NY-ESO-1-specific TCR. CD11b-deficient cells exhibited reduced killing of NY-ESO-1-expressing tumor cells compared with CD11b-sufficient T cell controls (**Fig. 5c–d and Extended Data Fig. 5b–c)**. Phospho-kinase profiling further showed differences in several signaling proteins following TCR activation **(Extended Data Fig. 5d–e)**

To further explore the translational relevance of CD11b in adoptive T cell therapy, we extended our analysis to human T cells expressing a second-generation CD19-targeting CAR containing 4-1BB and CD3z signaling domains (19BBz)^24, 25^. After in vitro expansion, a substantial fraction of 19BBz CAR-T cells expressed CD11b at frequencies comparable to those observed in UTD control T cells (**Fig. 5e),** indicating that CAR expression does not intrinsically alter CD11b surface expression.

To determine whether CD11b enriched for CAR-T cells with greater cytotoxic potential, we FACS-purified CD11b⁺ and CD11b^−^ 19BBz CAR-T cells and assessed their antitumor activity against SK-MEL-28 melanoma cells engineered to express a truncated variant of the CD19 receptor (SK-MEL-28 CD19t). Consistent with our observations in antigen-specific CD8^+^ T cells, CD11b⁺ 19BBz CAR-T cells mediated more efficient target-cell killing than their CD11b^−^ counterparts **(Fig. 5f and Extended Data Fig. 5f).** Together, these findings identify CD11b as a candidate surface marker for enriching TCR- and CAR-T cell products with enhanced cytotoxic activity.

## Discussion

Tumor-infiltrating CD8⁺ T cells comprise heterogeneous populations that differ in antigen specificity, differentiation state, and effector potential. Surface markers such as PD-1, CD39, and CD103 have been instrumental in enriching for tumor-reactive T cells across human solid tumors^3, 4^, but they do not fully resolve cytotoxic competence within the intratumoral CD8⁺ T cell compartment. This distinction is particularly relevant in solid tumors, which contain abundant bystander CD8⁺ T cells and tumor-reactive populations themselves encompass functionally diverse states^1, 26, 27^. Recent multimodal analyses further support this concept, showing that effector-potent CD8⁺ T cells can be delineated within exhausted intratumoral populations by integrating transcriptional and protein-level activation states^28^. Here, we identified CD11b as a TCR-inducible surface integrin that marks cytotoxic competence within tumor-reactive CTLs and contributes to their effector function. Within antigen-specific CD8⁺ T cell populations, CD11b distinguished cells with enhanced degranulation and inflammatory cytokine production. CD11b therefore adds a functional layer to established tumor-reactivity markers by identifying a cytotoxic-competent state within the broader antigen-experienced CD8⁺ T cell pool.

CD11b is best known as the αM chain of the CD11b/CD18 (αMβ2) integrin complex expressed by myeloid cells, where it regulates adhesion, migration and innate immune responses^29^. In tumors, CD11b signaling has also been implicated in the regulation of myeloid cell polarization, and pharmacological CD11b agonism can promote antitumor immunity^30, 31, 32^. Earlier studies in infection and autoimmunity models reported CD11b expression on subsets of effector CD8⁺ T cells^33, 34, 35, 36^, but its regulation and functional relevance in tumor-reactive CTLs remained unclear. Our data show that *ITGAM* is transcriptionally induced following TCR stimulation in naive CD8⁺ T cells as part of an inducible β2-integrin remodeling program triggered during antigen-driven T cell differentiation. The observation that CD11b is not uniformly expressed across antigen-specific CD8⁺ T cells further suggests that it marks a functional state rather than a specific T cell lineage. This heterogeneity may reflect differences in TCR signaling history, antigen persistence, or local tumor-derived cues, consistent with the view that intratumoral CD8⁺ T cell states are dynamically regulated rather than terminally fixed^26, 27^. Although CD11b⁺ CD8⁺ T cells were validated across multiple tumor models and detected in human samples, the prevalence and regulatory mechanisms governing CD11b induction across diverse tumor types and therapeutic settings remain to be defined. While our data support CD8^+^ T cell-intrinsic transcriptional induction following TCR activation, whether membrane exchange mechanisms such as trogocytosis^15, 43^ contribute to CD11b surface abundance in vivo remains to be determined.

This integrin remodeling has functional consequences at the immune synapse. CD11b localizes to the pSMAC, placing it within the adhesive and signaling architecture that coordinates TCR engagement, contact stability, and polarized secretion^7, 8^. LFA-1, the canonical CD11a/CD18 lymphocyte integrin, is constitutively expressed in T cells and has well-established roles in immune synapse formation and cytolytic granule release^9, 37^. More recently, magnesium sensing through LFA-1 was shown to promote its active conformation and enhance CD8⁺ T cell signaling, metabolic fitness, immune synapse formation, and antigen-specific cytotoxicity^18^. Our findings extend this principle by showing that inducible changes in β2-integrin composition can also regulate cytotoxic output. They support a model in which activation-induced CD11b is recruited as a non-canonical integrin to the pSMAC and contributes to cytotoxic effector function. The relevant ligand interactions, downstream mechanisms governing synaptic organization and granule release, and the extent to which these functions differ from those of LFA-1 remain unresolved. Together with emerging evidence that noncanonical integrin configurations can potentiate TCR signaling and antigen-specific cytotoxicity^11^, our results support a broader view of integrin composition as a dynamic determinant of cytotoxic T cell function.

Our findings also suggest that integrin composition may be exploitable in adoptive T cell therapy. CD11b expression promoted cytotoxic-granule release and identified CAR-T cells with enhanced killing capacity. Whereas pharmacological CD11b agonism have been studied predominantly in the myeloid compartment^30, 31, 32^, our results identify activated cytotoxic T cells as an additional CD11b-expressing population. In solid tumors, CAR-T cell adhesion and cytotoxicity depend on tumor-cell IFNγR–ICAM-1 signaling and CAR-T cell β2 integrins^38^. Integrin composition may therefore represent an additional cell-intrinsic feature amenable to engineering, alongside programs regulating T-cell fitness, persistence, resistance to exhaustion and effector function^39, 40, 41, 42^. Establishing whether CD11b can be used prospectively to select or engineer superior therapeutic T cell products will require validation across additional CAR designs, tumor contexts, and in vivo models. Together, our study identifies CD11b as an inducible β2-integrin component of cytotoxic CD8⁺ T cells that marks and enhances antitumor effector function. By integrating surface proteomic discovery with immune synapse organization and functional perturbations, we reveal an unexpected layer of integrin remodeling in tumor-reactive CD8^+^ T cells. More broadly, our data position integrin composition as a dynamic determinant of cytotoxic potency and a potential avenue for improving T cell-based cancer immunotherapies.

## METHODS

### Human samples

Primary human endometrial cancer specimens were collected and processed at the Vall d’Hebron Institute of Oncology (VHIO), Barcelona, Spain, and kindly provided by Dr. Alena Gros. The collection and use of these specimens were approved by the Vall d’Hebron Hospital ethical committee under protocol PR(AG)482/2017 and were conducted in accordance with the Declaration of Helsinki. All participants provided written informed consent.

De-identified, dissociated tumor cells from lung cancer, breast invasive ductal carcinoma, and breast invasive mammary carcinoma from consented patients were purchased from Discovery Life Sciences (Huntsville, AL).

### Mice

C57BL/6 were obtained from Charles River. Tg(TcraTcrb)1100Mjb/J (OT-I) mice (The Jackson Laboratory, JAX ID:003831) were used as a source of ovalbumin-specific CD8^+^ T cells. Female C57BL/6 mice were used for EO771 mammary and ID8-OVA ovarian tumor models. Female and male C57BL/6 mice were used for YUMMER1.7 and B16-OVA melanoma experiments, each experiment was sex-controlled, and no sex-dependent differences were observed. All mice were used at 8–12 weeks of age and housed under specific-pathogen free conditions in individually ventilated cages at the CIC bioGUNE Animal Facility at 21-22 °C and 39-50% relative humidity on a 12-h light/dark cycle. No formal statistical methods were used to predetermine sample size, and investigators were not blinded to mouse groups. All procedures were conducted in accordance to AAALAC International standards and approved by the Biosafety and Welfare Committee of CIC bioGUNE and the Committee on Bioethics and Animal Welfare and the Government of Bizkaia under licenses P-CBG-CBBA-0319, P-CBG-CBBA-0521, P-CBG-CBBA-1022, and P-CBG-CBBA-0225.

### Orthotopic breast tumor model

For the EO771 and EO771-OVA tumor models, 0.25×10^6^ EO771-GFP/Luc or EO771-OVA-GFP/Luc cells resuspended in 100 μl of incomplete RPMI-1640 medium were injected into the fourth mammary pads of anaesthetized C57BL/6 mice. A small skin incision was made adjacent to the fourth inguinal mammary gland, and the mammary fat pad was gently exposed and injected directly under visual control, using a surgical approach adapted from previously described methods^45^. Carprofen (10 mg/kg body weight) was administered subcutaneously (s.c.) 15 min before surgery and once daily for 3 days thereafter.

Tumors were measured with digital calipers every 2–3 days. Volume was calculated as: volume (mm^3^) = (π × length× width^2^)/6. Mice were euthanized when tumor volume exceeded 1500 mm^3^, or when any of the following prespecified endpoints was reached: >10% body weight loss, skin ulceration, or lethargy.

### Melanoma tumor models

For the YUMMER1.7 tumor model, 1×10^6^ cells were resuspended in 100 μl of incomplete DMEM/F12 and subcutaneously injected into the right flank of anaesthetized C57BL/6 mice. For the B16-OVA tumor model, 5×10^5^ cells were injected subcutaneously in the right flank of anaesthetized C57BL/6 mice. After tumors became palpable, tumor growth was monitored every 2–3 days and volume was calculated as described above. Humane endpoints were the same as those described for EO771 breast tumors.

### Ovarian cancer model

For ID8-OVA ovarian tumor model, 5×10^6^ cells were intraperitoneally (i.p.) injected in 100 μl of incomplete RPMI-1640 medium into anaesthetized C57BL/6 mice. Mice were monitored for body weight, posture, abdominal circumference and lethargy. Prespecified endpoints included hunched posture, lethargy or more than 20% increase in abdominal circumference or body weight attributable to intraperitoneal tumor growth.

### Mouse tissue processing

Mammary and melanoma tumors were excised at indicated time points, minced with razor blades in a cell culture dish and digested for 30 min at 37 °C in DMEM containing 5% FBS, collagenase IV (1 mg/mL; Sigma-Aldrich, #C0130), and DNase I (0.1 mg/mL; Merck, #10104159001), with vortexing every 10 min. Sample digests were passed through a 70 μm cell strainer and centrifuged at 450g for 5 min at 4 °C. Red blood cells were lysed for 2 min at room temperature with ACK lysis buffer (Gibco, A1049201). Cells were washed and resuspended in RPMI-1640 containing 10% FBS for downstream staining. Splenocytes were obtained by mechanical dissociation through a 70 μm cell strainer followed by ACK lysis buffer. Blood was collected from the submandibular vein into EDTA-coated tubes at indicated time points, and erythrocytes were lysed by ACK before flow cytometry. For mice bearing ID8-OVA tumors, the peritoneal cavity was lavaged with 5 mL of ice-cold PBS. The abdomen was gently massaged, lavage fluid was recovered after incision, and cells were centrifuged and processed for flow cytometry.

### Cell lines and culture conditions

EO771 mouse mammary adenocarcinoma cells (ATCC, #CRL-3461) were maintained in DMEM (Gibco, #41966-029) supplemented with 10% heat-inactivated fetal bovine serum (HI-FBS), 20 mM HEPES (Gibco, #15630056), and 1% penicillin/streptomycin. ID8 mouse ovarian epithelial cells (Merck, #SCC145) were maintained in RPMI-1640 medium (Gibco, #61870044) containing 10% H-FBS and penicillin/streptomycin. YUMMER1.7 H2B-GFP mouse melanoma cells (Merck, #SCC245) were maintained in DMEM/F12 medium (Gibco, #11320033) supplemented with 10% HI-FBS, MEM non-essential amino acids (Gibco, #11140035), and 1% penicillin/streptomycin. B16-F10 cells (ATCC, #CRL-6475) were cultured in DMEM containing 10% HI-FBS, 50 μM of β-mercaptoethanol (Thermo Fisher Scientific, #31350010) and 1% penicillin/streptomycin. SK-MEL-28 human melanoma cells (ATCC, HTB-72) were maintained in MEM containing 10% HI-FBS and 1% penicillin-streptomycin. Lenti-X 293T packaging cells (Takara Bio, #632180) were maintained in DMEM supplemented with 10% HI-FBS and 1% penicillin/streptomycin. All cell lines were cultured at 37 °C with 5% CO2 and used during exponential growth for implantation or functional assays. All lines routinely tested negative for mycoplasma. Commercial cell lines were not independently authenticated after acquisition. Engineered variants were verified for the introduced antigen or reporter by flow cytometry and/or fluorescence imaging and were monitored for expected morphology and growth. No line was listed in the International Cell Line Authentication Committee registry (v.11).

### Generation of engineered tumor cell lines

Unless otherwise stated, tumor cell lines were transduced with a lentiviral luciferase-EGFP (GFP/Luc) reporter, and GFP-positive cells were sorted by FACS. To generate OVA-expressing targets, the chicken ovalbumin insert from pCl-neo-cOVA (gift from Maria Castro; Addgene plasmid #25097) was cloned into pLV-MSCV-puromycin (Genscript). At 48 h after transduction, EO771-OVA, ID8-OVA and B16-OVA cells were selected with puromycin (1 μg/mL; Thermo Fisher Scientific, #A1113803), with replacement of selection medium every 3 days for 1 week.

To generate CD19-expressing targets, a truncated human CD19 sequence lacking the intracellular domain (CD19t)^46^ was synthesized and cloned into pLV-MSCV (Genscript). SK-MEL-28-CD19t cells were generated by lentiviral transduction followed by FACS isolation of CD19-positive cells.

To generate NY-ESO-1-expressing targets, NY-ESO-1 and HLA-A*02:01 coding sequences were synthesized and cloned (Genscript) into pLV-MSCV-puromycin and pLV-MSCV-blasticidin, respectively. SK-MEL-28-NY-ESO-1 cells were generated by co-transduction with the NY-ESO-1 and HLA-A*02:01 vectors. At 48 h, cells were selected with puromycin (1 μg/mL) and blasticidin (10 μg/mL; Thermo Fisher Scientific, #A1113903), with replacement of selection medium every 3 days for 1 week.

### Isolation and activation of human T cells

Human primary T cells were obtained from Buffy coats of healthy donors (Biobanco Vasco, BIOEF) after appropriate approval of the Ethical and Scientific Committees (code CEIC E19-75). Briefly, peripheral blood mononuclear cells (PBMCs) were isolated by density gradient centrifugation using SepMate-50 columns (STEMCELL Technologies, #85450) with Ficoll-Paque PLUS (Cytiva, #17144003) as per manufacturer’s instructions. Naive CD8^+^ T cells, total CD8^+^ T cells and total CD3^+^ T cells were isolated by negative selection using the EasySep™ Human Naïve CD8^+^ T Cell Isolation Kit (STEMCELL Technologies, 19258), EasySep™ Human CD8^+^ T Cell Isolation Kit (STEMCELL Technologies, 17953) and Human T Cell Enrichment Kit (STEMCELL Technologies, 19051), respectively, following manufacturer’s instructions. Purity was routinely confirmed by flow cytometry to be >95%.

Unless otherwise stated, T cells were activated with anti-CD3/CD28 Dynabeads (Thermo Fisher Scientific, #11132D) at a bead-to-cell ratio of 1:1 in CST OpTimizer T Cell Expansion medium (Thermo Fisher Scientific, #A1048501) containing 5% CTS Immune Cell Serum Replacement (Thermo Fisher Scientific, #A2596101) and recombinant human IL-2 (100 IU/mL; Miltenyi Biotec, #130-097-748). Beads were magnetically removed 4-6 days after activation, and cultures were replenished with fresh medium every 2 days.

For longitudinal analysis of integrin expression, purified naive human CD8^+^ T cells were stimulated in plates coated with agonistic anti-CD3 (5 μg/mL; OKT3, BD Biosciences, #567118) and soluble anti-CD28 (5 μg/mL; CD28.2, eBiosciences, #16-0289-38) monoclonal antibodies in complete OpTimizer medium. Activated CD8^+^ T cells were collected at indicated time points for flow cytometry or RT-qPCR.

### Isolation and activation of mouse CD8^+^ T cells

Spleens from C57BL/6 or OT-I mice were mechanically dissociated and filtered through a 70-μm cell strainer. Total or naive CD8^+^ T cells were isolated by negative selection using the EasySep Mouse CD8^+^ T Cell Isolation Kit (StemCell Technologies, 19853) or EasySep™ Mouse Naïve CD8+ T Cell Isolation Kit (StemCell Technologies, 19858). For CD8^+^ T cell activation, 1×10^6^ purified CD8^+^ T cells were stimulated with anti-CD3/CD28 Dynabeads (Thermo Fisher Scientific, #11452D) at a bead-to-cell ratio of 1:1 in RPMI-1640 containing 10% HI-FBS, 20 mM HEPES (Gibco), 1 mM sodium pyruvate (Gibco), 50 μM β-mercaptoethanol, MEM non-essential amino acids (Gibco), recombinant human IL-2 (100 IU/mL; Miltenyi Biotec), and 1% penicillin/streptomycin). Fresh medium was added every two days, and cells were used at indicated time points post-activation.

### Lentiviral expression constructs

The second-generation anti-CD19 CAR (19BBz) used was previously described^42^ and contained an FMC63-derived single-chain variable fragment (scFv)^47^, a human CD8α hinge and transmembrane region, a human 4-1BB costimulatory domain, and a human CD3z signaling domain. The CAR coding sequence was linked by a self-cleaving P2A peptide to a truncated nerve growth factor receptor (tNGFR), which served as transduction and selection marker.

The NY-ESO-1-specific TCR was based on the affinity-enhanced 1G4 α95:LY TCR, which recognizes NY-ESO-1 residues 157-165 (SLLMWITQC) presented by HLA-A*02:01 and has a reported KD=730 nM^48^. The TCR coding sequence was synthesized and cloned downstream of the EF1α promoter in a third-generation lentiviral transfer vector (Genscript). Human TRAC and TRBC constant regions incorporated the T48C and S57C substitutions, respectively, which introduce an additional interchain disulfide bond to favor pairing of the introduced TCRα and TCRβ chains^49^. The TCR cassette was linked to tNGFR by a self-cleaving T2A peptide, which was used as transduction and selection marker.

The coding sequence of full-length human CD11b (ITGAM; UniProt accession P11215) was synthesized and cloned into the pLV-MSCV lentiviral transfer vector (GenScript). The CD11b I332G variant was generated by introducing an Ile332Gly point substitution into the wild-type CD11b coding sequence (GenScript). For selection purposes, the CD11b coding sequence was linked via a self-cleaving P2A peptide to a blasticidin-resistance gene.

For some fluorescence-imaging experiments, the coding sequence of the αTAG–mEGFP reporter was synthesized and fused in frame to the C terminus of wild-type CD11b (Genscript), generating CD11b–αTAG–mEGFP.

### Lentiviral production

To generate lentiviral particles, 5×10^6^ Lenti-X 293T cells were seeded in 100 mm dishes. After 24 h, cells were transfected with 5 µg transfer plasmid, 4 µg psPAX2 (Addgene, 12260) and 1.5 µg VSV-G (Addgene, 8454) using jetPEI (Polyplus, 101-10N) according to manufacturer’s instructions. Supernatants were collected 48 h later, lentiviral particles were harvested from the supernatant, 0.45 μm filtered, and incubated with LentiX Concentrator (Takara, 631232) as per manufacturer’s protocol at 4 °C overnight. Lentiviral particles were then concentrated by centrifugation at 1500g for 45 min, aliquoted, and stored at −80 °C until use.

### Generation of human CAR-T cells

Human CD3^+^ T cells were activated with anti-CD3/CD28 Dynabeads as described above. On day 2 after activation, cells were transduced with 19BBz lentivirus at multiplicity of infection (MOI) 6 in the presence of polybrene (8 μg/mL; Merck, TR-1003) by spinoculation at 800g for 1 hour at 32°C. Cells were maintained at 0.5 × 10^6^ cells/mL in complete CTS OpTmizer medium containing IL-2, with fresh medium added every 2–3 days. On day six after activation, beads were removed and transduced cells were enriched using the EasySep Human CD271 Positive Selection Kit II (STEMCELL Technologies, 17849). CAR-T cells were expanded for 10–12 days, cryopreserved in FBS containing 10% DMSO and stored in liquid nitrogen. CAR-T purity was routinely >85% by flow cytometry.

### Generation of human NY-ESO-1 TCR-T cells

Activated human CD8^+^ T cells were transduced on day 2 with NY-ESO-1 TCR lentivirus at MOI 6 in the presence of polybrene and spinoculated as described above. On day 6 after activation, NGFR-positive cells were enriched with the EasySep Human CD271 Positive Selection Kit (STEMCELL Technologies, 17849) and expanded by adding fresh medium every 2–3 days. Cells were expanded for 10–12 days, and transduction efficiency was routinely >90% based on NGFR expression by flow cytometry.

### CD11b overexpression

For untagged wild-type or I332G CD11b overexpression, activated human CD3^+^ T cells were transduced on day 2 at MOI 3 as described above. Blasticidin selection (10 μg/mL) was initiated 48 h after transduction and continued for 1 week. CD11b overexpression was confirmed by flow cytometry. For the indicated imaging experiments, human CD8^+^ T cells were transduced on day 2 with lentivirus encoding αTAG-mEGFP-tagged wild-type CD11b at MOI 6. Cells were expanded for 10–12 days, and preparations containing >30% mEGFP-positive cells were used.

### CRISPR/Cas9 editing of human T cells

Isolated human CD8^+^ T cells were activated as described above. CD8^+^ T cells were electroporated using the Neon™ Transfection System (Thermo Fisher Scientific) according to manufacturer’s instructions. Briefly, 2×10^5^ activated CD8^+^ T cells were debeaded at day 4 post-activation, resuspended in 5 μL of buffer R and combined with 5 μL of RNP solution. Unless otherwise indicated, sgRNAs were selected from predesigned CRISPR-Cas9 Trueguide RNAs from Thermo Fisher. Per reaction, 6 pmol of sgRNA were mixed with 1 μg of TrueCut™ Cas9 Protein v2 (Thermo Fisher Scientific, A36498) followed by incubation at room temperature for 30 min. Chemically synthesized sgRNA targeting *ITGAM* exon 5 (sequence: GCCGTAGGTTGGATCCAAAC), *ITGAL* exon 3 (sequence: TGCCCGACTGGCACTGATAG)^18^, or *ITGB2* exon 5 (sequence: CAGCGTTCAACGTGACCTTC) were used. Cells were electroporated with buffer alone as negative control (KO control). An appropriate nucleofector program was applied (1600 V, 10 ms pulse width, and 3 pulses). Editing was assessed by loss of CD11b, CD11a or CD18 surface expression by flow cytometry.

For the generation of CD11b-deficient NY-ESO-1 TCR-specific CD8^+^ T cells, NY-ESO-1 TCR T cells were FACS sorted based on CD11b expression, reactivated for 4 days with anti-CD3/CD28 Dynabeads, electroporated with buffer alone (CD11b^+^ ctrl) or CD11b-targeting RNPs, and expanded for 6 days before functional assays.

### Flow cytometry and cell sorting

Single-cell suspensions from indicated tissues were prepared as described above. DAPI (Invitrogen; 1:20,000) or LIVE/DEAD Fixable Blue (Thermo Fisher Scientific; 1:1000) was used to exclude non-viable cells. Samples were washed in cytometry staining buffer (Thermo Fisher Scientific) and incubated for 30 min at 4 °C with fluorescently conjugated antibodies. The human or mouse TruStain FcX antibody was used to prevent non-specific binding. See Supplementary Table 1 for antibodies used for flow cytometry.

Mouse antigen-experienced CD8^+^ T cells were defined as live singlet CD45^+^TCRβ^+^CD8α^+^CD44^+^ cells. Mouse naive CD8^+^ T cells (CD8^+^ TN) were defined as live singlet CD45^+^TCRβ^+^CD8α^+^CD44^−^CD62L^+^ cells. Human CD8^+^ tumor infiltrating lymphocytes were defined as live singlet CD45^+^CD3^+^CD56^−^CD8^+^ cells. Full gating strategies are shown in Extended Data Fig. 2 a,f.

Data were acquired on a four-laser BD FACSymphony (BD Bioscience) and analyzed in FlowJo v10.10.1. Fluorescence-minus-one controls were used to define CD11b expression and other low-frequency or continuous populations where required. Cell sorting was performed on BD FACSAria Fusion (BD Bioscience), and post-sort purity and viability were confirmed to be >95% by flow cytometry.

### LEGENDScreen cell surface proteome profiling

Primary EO771 tumors (n = 4 mice), spleens from tumor-bearing mice (n = 4 mice), spleens from tumor-free mice (n = 4 mice) and mammary fat pads from tumor-free mice (n = 10 mice) were collected and processed as described above. Single cell suspensions were pooled within each tissue category before staining, yielding one pooled sample per category. Cells were stained with LIVE/DEAD^TM^ Fixable Blue for 30 min at 4 °C, washed, adjusted to 1 × 10^7^ cells/mL and incubated with mouse TruStain FcX Fc-receptor blocking reagent (1:50, Biolegend) for 10 min at room temperature. Samples were then barcoded according to tissue origin using distinct anti-CD45-fluorochrome conjugates (tumor, APC; tumor-bearing mice spleen, BUV496; tumor-free mice MFP, APC+BUV496; tumor-free mice spleen, BUV805). After staining for 30 min at 4 °C, the samples were washed, combined, and stained for 30 min at 4 °C with a backbone panel comprising the following antibodies: CD8a-BUV395, CD4-BV786, TCRβ-BV480, TCRγδ-R718, NK1.1-APC/F750, CD44-AF488, CD62L-BV650, PD-1-BV605, CD39-PE/Dazzle594, CD103-BV711, CD69-PE/Cy7, CD127-BV421 and CD25-PerCP-eF710. Cells were washed, resuspended at 4 × 10^6^ cells/mL, and subjected to the LEGENDScreen Mouse PE Kit (Biolegend, #700009) containing 255 PE-conjugated monoclonal antibodies against various mouse cell surface markers, plus 11 mouse, rat, and hamster Ig isotype controls, arrayed on three 96-well plates. Staining, washing and fixation were performed according to the manufacturer’s protocol, and samples were acquired on a four-laser BD FACSymphony (BD Biosciences).

Flow-cytometry data were analyzed using FlowJo v10. After the exclusion of debris, doublets and dead cells, the four pooled samples were deconvoluted according to their CD45 barcode, and pre-gated on antigen-experienced CD44^+^ CD8^+^ TCRβ^+^ T cells. Marker-positive gates in the PE channel were defined using the corresponding internal isotype controls, and markers detected in ≥10% of cells in at least one tissue were retained for analysis. For each marker, tissue enrichment was calculated as the difference between the frequency of marker-positive cells in each tissue and the mean frequency across the remaining three tissues. Tissue-enrichment scores were visualized as a heatmap, with markers ordered by decreasing tumor-enrichment score.

### Ex vivo restimulation and intracellular cytokine staining

Tumor single-cell suspensions (2×10^6^ cells) were stimulated at 37 ^°^C for 16 h with eBioscience Cell Stimulation Cocktail plus protein transport inhibitors (Thermo Fisher Scientific, 00-4975-03) along with anti-CD107a antibody. After staining for viability with LIVE/DEAD^TM^ Fixable Blue for 30 min at 4 ^°^C, Fc blocking and staining for cell-surface markers for identification of antigen-experienced T cells (anti-mouse CD45, CD8, CD4, TCRβ, CD44, PD-1 and CD11b). Cells were then fixed and permeabilized using the IntraStain kit (Agilent, K231111-2) and stained with antibodies against TNF, IFNγ, granzyme B and perforin as per manufacturer’s instructions. T cell staining was analyzed by flow cytometry analysis as described above.

### CD11b molecular conformation

To evaluate CD11b molecular conformation changes induced by divalent cations, human CD8^+^ T cells overexpressing wild-type CD11b or the constitutively active CD11b I332G variant were expanded for 10 days. Cells were incubated for 1 h in the presence or absence of 3 mM MnCl2. The activation state of CD11b can be assessed using monoclonal antibodies recognizing activation-dependent conformational epitopes^50^. The active αI-domain high-affinity conformation of CD11b was detected by flow cytometry using clone CBRM1/5, whereas total CD11b was measured using conformation-independent clone M1/70.

### CD11b^+^ CD8^+^ tumor infiltrating T cell immunofluorescence

For confirmation of single-cell expression of αβTCR, CD8 and CD11b, fresh EO771 tumors were dissociated, and cells were stained as described in the flow cytometry section with CD11b-BV421, CD8-FITC and TCRβ-APC antibodies. Live singlet CD11b^+^TCRβ^+^CD8^+^ T cells were isolated by FACS, mounted immediately in ProLong Gold antifade medium (Thermo Fisher Scientific, P10144) and imaged on a Leica SP8 Lightning confocal microscope equipped with a 63× Plan Apochromat 1.4-NA objective.

### Preparation of glass-supported lipid bilayers

Acid- and plasma-cleaned (5 min) coverslips (Coverslip Glass D, 0.17 ± 0.005 mm; Schott Nexterion, 1472315) were mounted onto Sticky-Slide VI0.4 chambers (Ibidi, 80608) to assemble six-well imaging chambers. Glass-supported lipid bilayers (SLBs) were formed by incubating a liposome mixture containing 0.1875% vol/vol of 0.4 mM CapBio, 12.5% vol/vol 0.4 mM DGS-NTA(Ni) in a 0.4 mM DOPC matrix on the glass for 30 min at room temperature. Excess liposomes were removed by three washes with 1% bovine serum albumin-containing HEPES Buffered Saline (HBS/BSA) buffer. Then, SLBs were functionalized with 12-His-tagged mouse ICAM-1 (200 molecules per µm^2^) and anti-CD3ε Fab (clone UCHT1; 30 molecules per µm^2^). Recombinant protein densities were calibrated using bead-supported lipid bilayers as previously described^51,52^. Following incubation for 30 min at room temperature and three washes, T cells were stimulated on SLB (0.75×10^6^ cells/well) and incubated for 10 min at 37 °C. Cells were immediately fixed for 10 min with prewarmed 4% PFA in PBS containing 2 mM MgCl2, washed three times with HBS/HSA buffer, and either stained with anti-CD11b (AF647, clone M1/70, 1:100, BD Biosciences) to assess colocalization with mouse ICAM-1 (directly conjugated to AF405), or with antibodies against cytotoxic particles, including anti-human perforin (AF488, clone dG9, 1:100,BioLegend), and anti-human/mouse granzyme B (in-house conjugated AF568, clone GB11, 1:100, BioLegend). After overnight incubation, SLBs were washed four times with HBS/BSA buffer and imaged by total internal reflection fluorescence microscopy (TIRFM).

### TIRFM

We used an Olympus IX83 inverted microscope equipped with four diode LASERs providing illumination as four independent lines at 405 nm, 488 nm, 561 nm and 640-nm, an Olympus UApON 150×/1.45 NA objective, and a Photometrics Prime 95B back-illuminated sCMOS camera. Cell–SLB contacts were identified and focused using internal reflection microscopy (IRM) generated with LED illumination. Then, fluorescence images were acquired without previous LASER illumination and included IRM images. Images were analyzed using Fiji/ImageJ. Colocalization between CD11b and ICAM-1 within the synaptic contact area was quantified using the EzColocalization plugin with manual and Costes-assisted thresholding and expressed as Pearson’s correlation coefficient and the two directional Manders’ colocalization coefficients^53^. The latter indicates the total pixels of CD11b colocalizing with ICAM-1, and vice versa.

### Phospho-kinase array analysis

Isolated primary human CD8^+^ T cells were activated using anti-CD3/CD28 Dynabeads as described above. On day 6 after activation, CD11b-expressing CD8^+^ T cells were sorted and electroporated with buffer alone (CD11b^+^ ctrl) or ITGAM-targeting RNPs. After a further 6 days of culture, cells were restimulated with plate bound agonistic anti-CD3 antibody (2 µg/mL) for 6 h. Cell lysates were prepared in RIPA buffer and analyzed using the Proteome Profiler Human Phospho-Kinase Array Kit (R&D Systems; ARY003C) according to the manufacturer’s instructions. Chemiluminescent signals were detected using an iBright Imaging System (Thermo Fisher Scientific). Signal intensities were quantified and background-subtracted using ImageJ and normalized to the array reference controls. Duplicate technical spots for each analyte were averaged, and fold change was calculated relative to control-electroporated cells.

### Quantitative reverse-transcription PCR

Total RNA was extracted from human naive CD8^+^ T cells at indicated time points using the NucleoSpin RNA kit (Macherey-Nagel, 740955.250). The cDNA was synthesized from 1 μg of purified RNA using M-MLV reverse transcriptase and random primers (Thermo Fisher Scientific). Quantitative PCR reactions were performed in a Viia 7 Real-Time PCR system (Thermo Fisher Scientific) using PerfeCTa SYBR Green SuperMix reagent (Quantabio) and gene-specific primers. Relative expression levels were calculated as transcript levels of target genes relative to 18S housekeeping gene. Human primer sequences were: *ITGAM*: Fw, GGAACGCCATTGTCTGCTTTCG; Rv, ATGCTGAGGTCATCCTGGCAGA; *ITGAX*: Fw, GATGCTCAGAGATACTTCACGGC; Rv, CCACACCATCACTTCTGCGTTC; *ITGB2*: Fw, AGTCACCTACGACTCCTTCTGC; Rv, CAAACGACTGCTCCTGGATGCA; *ITGAL*: Fw, CTGCTTTTGCCAGCCTCTCTGT; Rv, GCTCACAGGTATCTGGCTATGG, and *18S* rRNA: Fw, CGGCGACGACCCATTCGAAC; Rv, GAATCGAACCCTGATTCCCCGTC.

### Bulk RNA-sequencing

Human CD8⁺ T cells were isolated from PBMCs from three healthy donors as described above. On day 12 after initial activation with anti-CD3/CD28 Dynabeads, cells were sorted into viable CD11b⁺ and CD11b⁻ populations and restimulated for 24 h with plate-bound agonistic anti-CD3 antibody (2 μg/ml). Total RNA was extracted using the NucleoSpin RNA kit (Macherey–Nagel, 740955.250). RNA concentration was measured using the Qubit RNA HS Assay Kit (Thermo Fisher Scientific, Q32855), and RNA integrity was assessed on an Agilent 2100 Bioanalyzer. All samples had RNA integrity number (RIN) values >9. Strand-specific, poly(A)-selected libraries were prepared from 250 ng total RNA using the TruSeq Stranded mRNA Library Prep Kit (Illumina, 20020594) according to the manufacturer’s instructions. Libraries were sequenced on an Illumina NovaSeq 6000 using 2 × 101-bp paired-end reads.

### RNA-seq data processing

Raw FASTQ files were quality-controlled using FastQC v0.11.6, with reports summarized using MultiQC v1.13. Adapter trimming was performed with fastp v0.23.2, and trimmed reads were aligned to the human hg38 reference genome using STAR v2.7.10b. PCR duplicates were removed using Picard v2.27.5. Processed BAM files were assessed using FastQC/MultiQC, samtools v1.16.1, and Picard v2.27.5. Gene-level read counts were generated from the processed BAM files using featureCounts v2.0.3. Differential expression between donor-matched CD11b⁺ and CD11b⁻ CD8^+^ T cells was analyzed from raw counts using *DESeq2,* with donor included as a blocking factor. P values were adjusted using the Benjamini–Hochberg method, and genes with an adjusted P value <0.05 were considered differentially expressed, without an additional fold-change threshold. For heatmap visualization, selected differentially expressed genes were plotted using DESeq2-normalized counts log2-transformed as log2(count + 1) and scaled across samples for each gene (row z-score).

### Gene ontology and gene set enrichment analysis

GO Biological Process over-representation analysis was performed using clusterProfiler on differentially expressed genes higher in CD11b⁺ cells, using genes entering differential-expression testing with defined adjusted P values and Entrez identifiers as the background. Significant terms were defined at Benjamini–Hochberg-adjusted P < 0.05. Redundancy among significant terms was reduced using Wang semantic similarity (simplify, cutoff = 0.7), after which representative non-redundant terms were selected for visualization.

Preranked gene set enrichment analysis was performed on the complete ranked transcriptome using the DESeq2 Wald statistic and published human CD8⁺ T-cell Cytotoxic and Cytokine signatures^44^. Benjamini–Hochberg correction was applied across tested signatures.

### Real-time cytotoxicity assays

Human melanoma SK-MEL-28-CD19t, SK-MEL-28-NY-ESO-1 or mouse B16-OVA cells were seeded at 0.25×10^5^ cells per well in a 96-well xCELLigence E-Plate (Agilent). 24 h later, human NY-ESO-1 TCR T cells, 19BBz CAR-T cells or mouse OT-I CD8^+^ T cells, were added at a 0.25:1 ratio and the target cell index was recorded every 10 min in the xCELLigence Real-Time Cell Analysis (RTCA) Analyzer (Agilent).

For OT-I CD8^+^ T cell cytotoxicity assays, purified CD8^+^ T cells were activated and expanded as described above. Following expansion, cells were stained with anti-mouse CD11b monoclonal antibody (clone M1/70, eBioscience) and sorted by FACS into CD11b⁺ and CD11b⁻ populations. Sorted CD11b⁺ and CD11b⁻ OT-I CD8^+^ T cells were recovered in complete mouse T cell media overnight, and subsequently used in cytotoxicity assays against OVA-expressing B16 target cells.

### Quantification and statistical analysis

Statistical analyses were performed using GraphPad Prism version 11. The experimental sample size and statistical test for each analysis are provided in the corresponding figure legend. Independent mice and independent human donors were treated as biological replicates; replicate wells from the same mouse, donor or cell preparation were treated as technical replicates and were averaged before inferential testing unless otherwise stated. All tests were two-sided, and data are presented as mean ± s.e.m unless otherwise indicated.

## Data availability

Transcriptomics data have been deposited in the Gene Expression Omnibus database under accession number GSE344884. The remaining data are available within the Article, Supplementary Information or Source Data file. Source data are provided with this paper.

## AUTHOR CONTRIBUTIONS

A.A.-V. conceived the study. A.A.-V. and J.E.-D. designed the experiments, performed most of the experimental work, and analyzed the data with assistance from A.B., E.P.-F., L.P.-G., P.V.-B., L.E.-M., I.O. and S.Y.L. A.M.A. provided technical support for the bulk RNA-sequencing experiments and contributed to data interpretation. K.Y. and P.F.C. provided intellectual input and designed and performed the immune synapse imaging experiments. M.M.C. provided intellectual input and assisted with the EO771 orthotopic breast tumor model. I.E.-S., M.F.S. and A.G. provided intellectual input and facilitated access to human samples. A.P. and J.E.-O. administered the project, supervised the study, provided resources and acquired funding. J.E.-D., J.E.-O., A.P. and A.A.-V., wrote the manuscript with input from all the authors.

### ACKNOWLEDGEMENTS

We thank the patients who donated tumor samples for this study and their families. We thank members of the Palazon and Ereño-Orbea laboratories for helpful discussions. We thank the Genome Analysis Platform at CIC bioGUNE for technical assistance. All illustrations were created with BioRender.com.

---

**Extended Data Figure 1.**
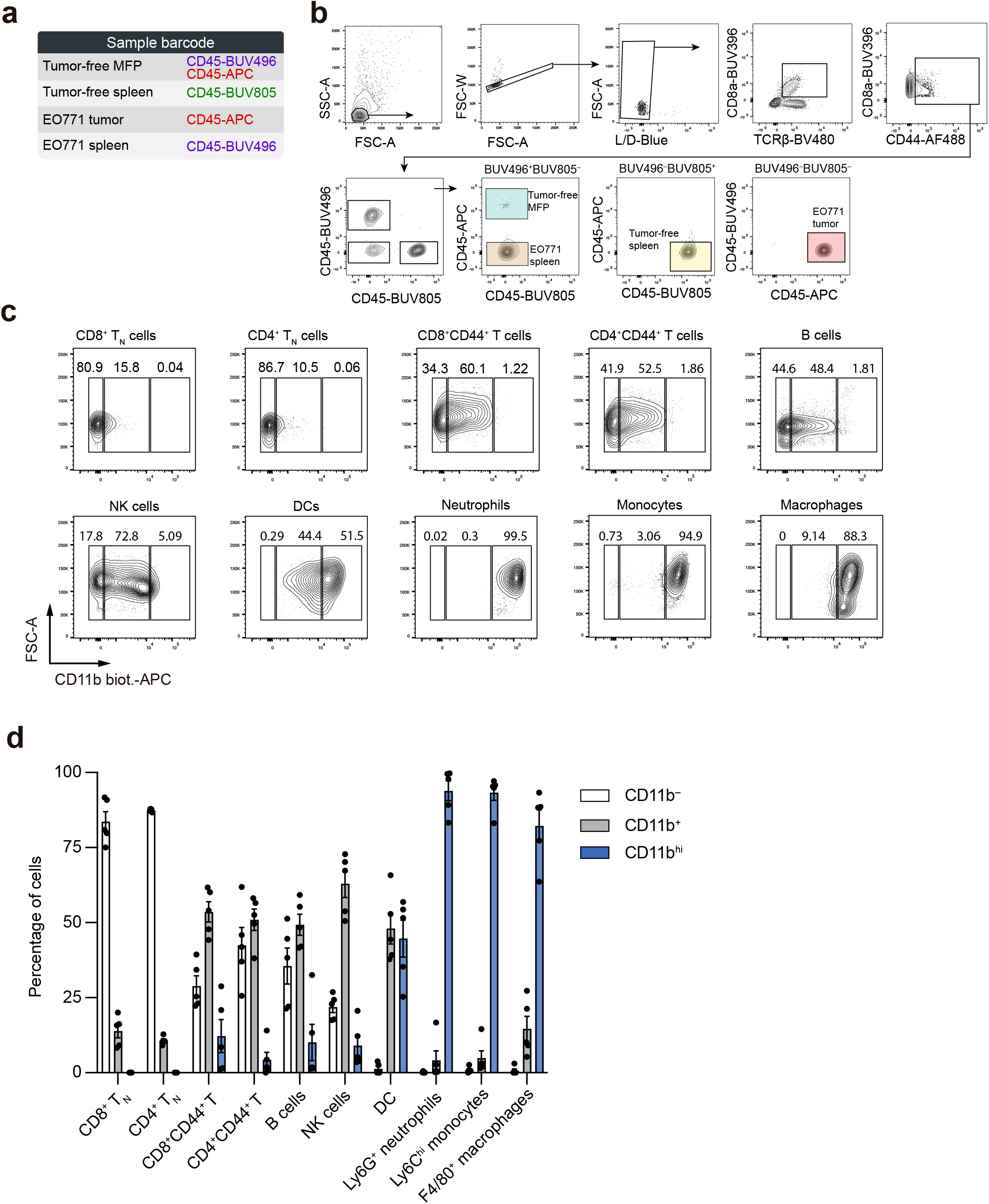
LEGENDScreen tissue barcoding and CD11b expression profiling. **a,** Tissue-barcoding strategy for tumor-free mammary fat pad (MFP), tumor-free spleen, EO771 tumor and spleen from EO771 tumor-bearing mice. **b,** Representative gating strategy for LEGENDScreen identification of antigen-experienced CD45⁺TCRβ⁺CD8α⁺CD44⁺ T cells and barcode deconvolution. **c,** Representative contour plots showing CD11b⁻, CD11b⁺, and CD11b^hi^ fractions within the indicated tumor-infiltrating immune cell populations. **d,** Quantification of the CD11b fractions shown in (**c**).

**Extended Data Figure 2.**
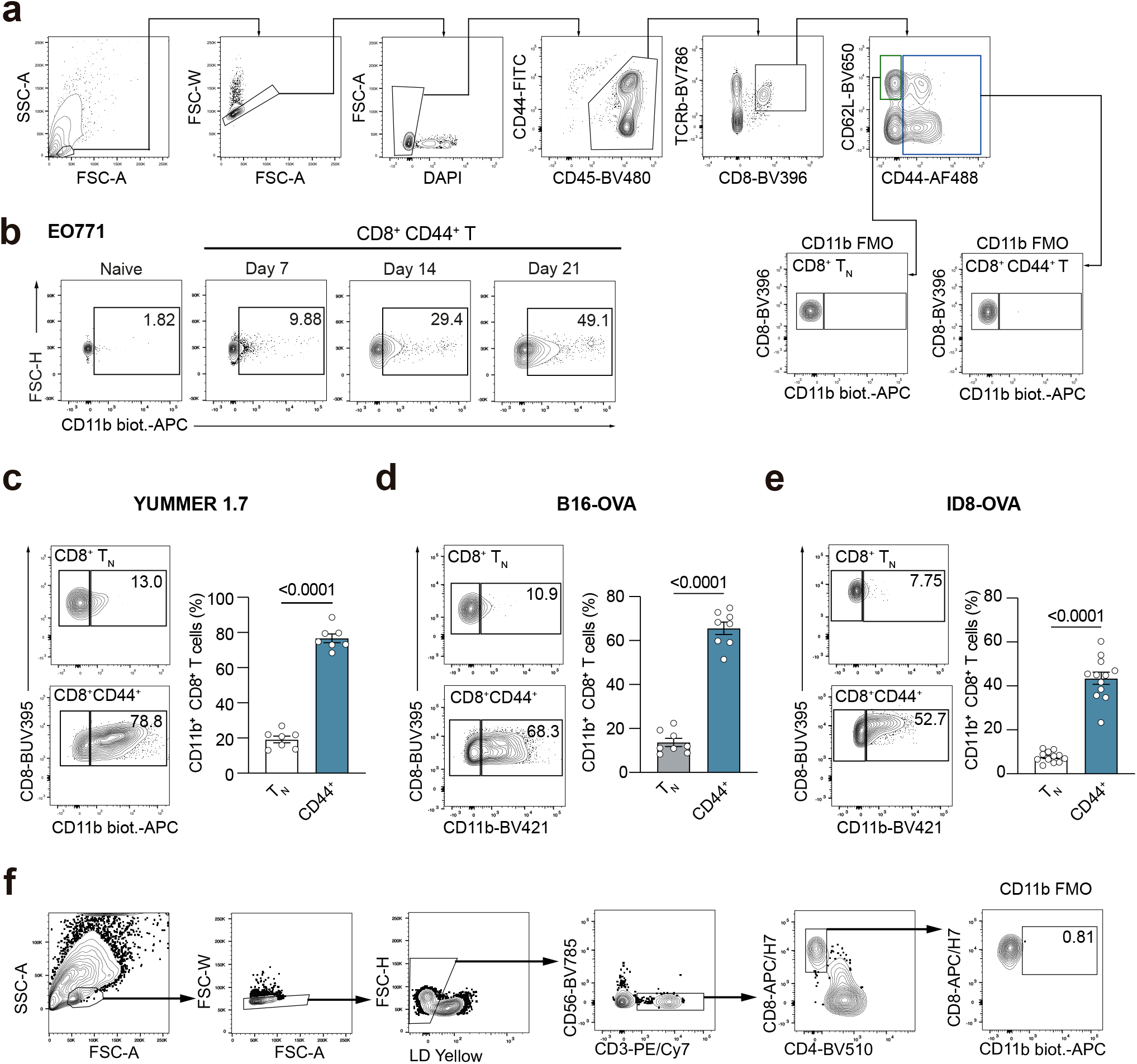
CD11b expression in CD8⁺ T cells across mouse and human tumors. **a**, Gating strategy used to identify mouse CD8⁺ T-cell populations. Fluorescence-minus-one (FMO) controls were used to define CD11b positivity. **b,** Representative contour plots of CD11b expression in naive CD8⁺ T cells and antigen-experienced EO771 CD8⁺ TILs on days 7, 14, and 21 after tumor implantation. **c–e,** Representative contour plots and quantification of CD11b expression in naive and antigen-experienced CD8⁺ T cells from YUMMER1.7 melanoma (day 14; n = 7 mice) (**c**), B16-OVA melanoma (day 19; n = 8 mice) (**d**) and ID8-OVA ovarian tumors (week 8; n = 12 mice) (**e**). CD8⁺ TN, CD62L⁺ CD44⁻ naive CD8⁺ T cells. **f,** Gating strategy used to identify human tumor-infiltrating CD8⁺ T cells. An FMO control was used to define CD11b positivity. P values were calculated using two-tailed, unpaired Student’s t test (**c–e**).

**Extended Data Figure 3.**
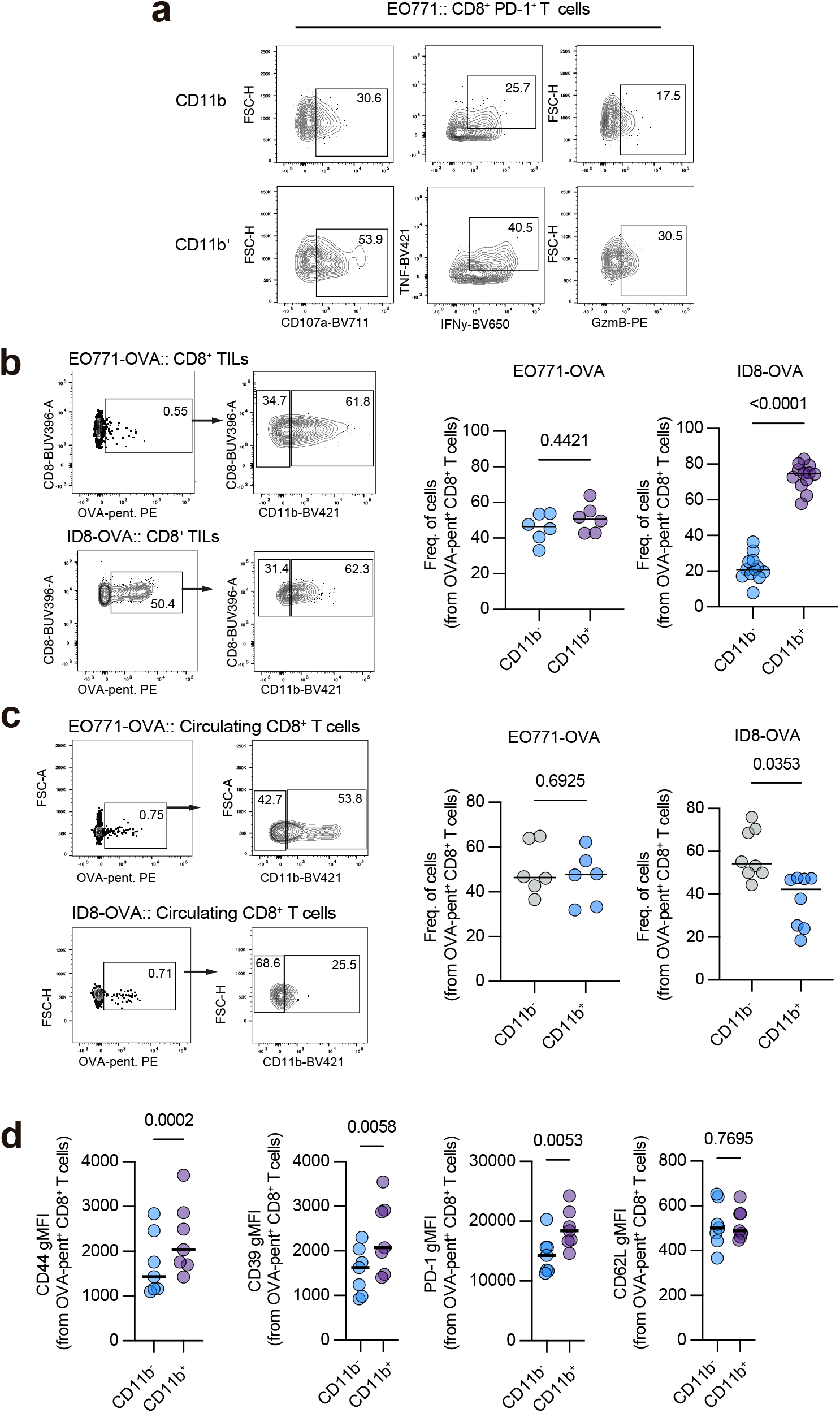
CD11b identifies highly functional tumor-reactive CD8⁺ T cells. **a,** Representative plots of CD107a^+^, IFN-γ^+^TNF^+^, and GzmB^+^ cells within CD11b⁻ or CD11b⁺ PD-1⁺ CD8⁺ TILs from EO771 tumors after overnight ex vivo stimulation. **b,** Representative plots and frequencies of CD11b⁺ cells among OVA-pentamer⁺ CD8⁺ TILs (**b**) and circulating CD8⁺ T cells (**c**) from EO771-OVA tumors (day 14; n = 6 mice) and ID8-OVA tumors (week 8; n = 12 mice). **d,** Phenotypic comparison of CD44, PD-1, CD39, and CD62L expression in CD11b⁺ or CD11b⁻ OVA-pentamer⁺ CD8⁺ TILs from B16-OVA tumors on day 21 (n = 7 mice). P values were calculated using two-tailed, paired Student’s t test (**b–d**).

**Extended Data Figure 4.**
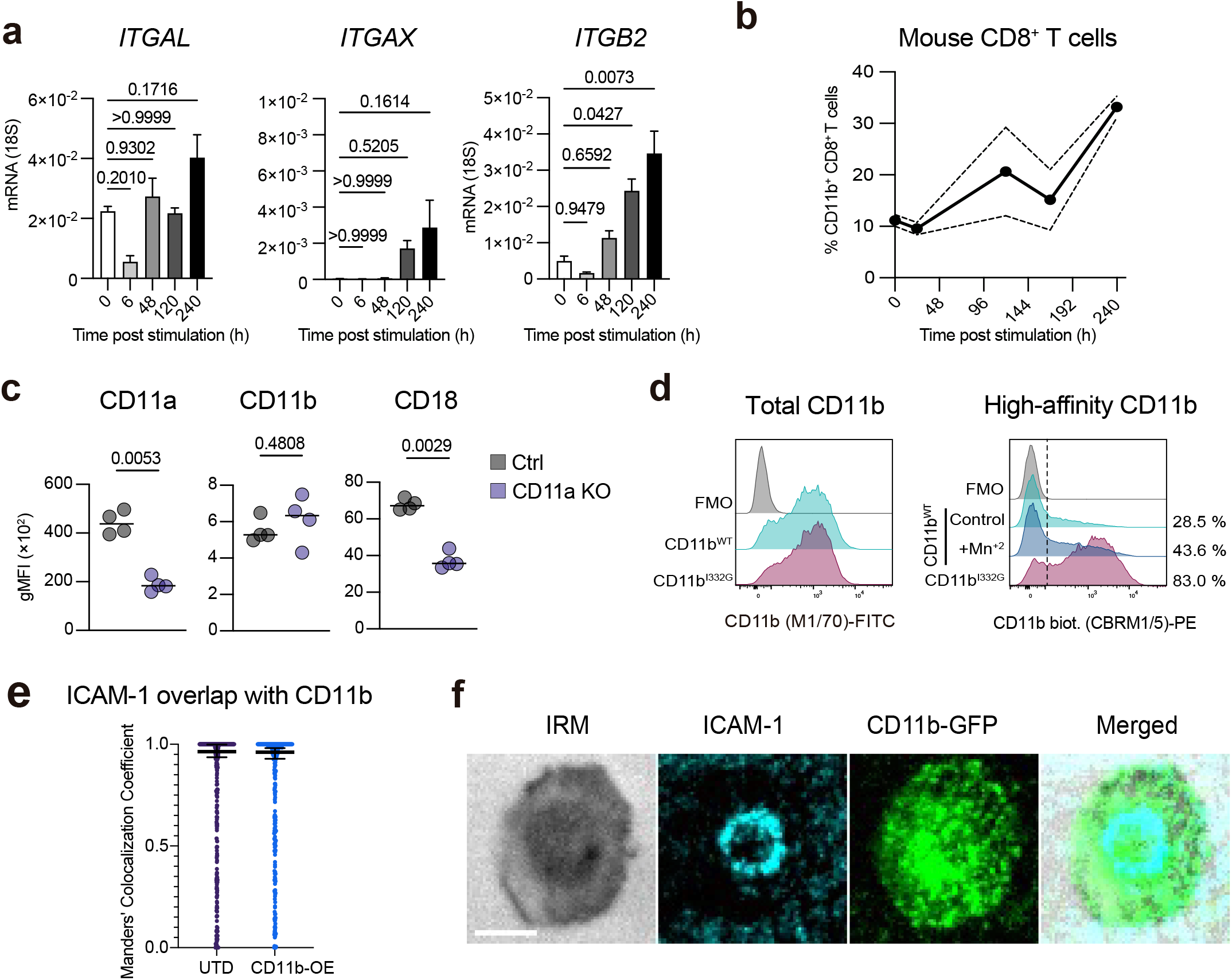
Activation-dependent β2-integrin expression and CD11b recruitment to the T cell immune synapse. **a,** *ITGAL*, *ITGAX*, and *ITGB2* transcript abundance in human CD8⁺ T cells at the indicated times after activation measured by qPCR (n = 3 donors). **b,** Frequency of CD11b^+^ mouse naive CD8^+^ T cells at the indicated times after in vitro activation (n = 3 mice). **c,** Surface abundance of CD11a, CD11b, and CD18 in control-electroporated (Ctrl) or CD11a-deficient human CD8⁺ T cells (n = 4 donors). **d,** Representative histograms of total CD11b expression (left) and high-affinity CD11b conformation (right) on human CD8⁺ T cells overexpressing wild-type CD11b (CD11b^WT^) or CD11b I332G, in the presence or absence of 3 mM MnCl2. Representative of three independent experiments. **e,** Representative TIRF microscopy images of immune synapses formed by CD11b–GFP-overexpressing human T cells on supported lipid bilayers (SLBs), showing the internal reflection microscopy (IRM) footprint, ICAM-1 (cyan), CD11b–GFP (yellow), and merged channels. Scale bar = 5 µm. **f,** Manders’ colocalization coefficients quantifying the fraction of ICAM-1 signal overlapping with CD11b within individual T cell–SLB contacts formed by untransduced (UTD) and CD11b-overexpressing (CD11b-OE) T cells. P values were calculated using one-way analysis of variance (ANOVA) with Tukey’s multiple-comparison test (**a**) and two-tailed, paired Student’s t test (**c**).

**Extended Data Figure 5.**
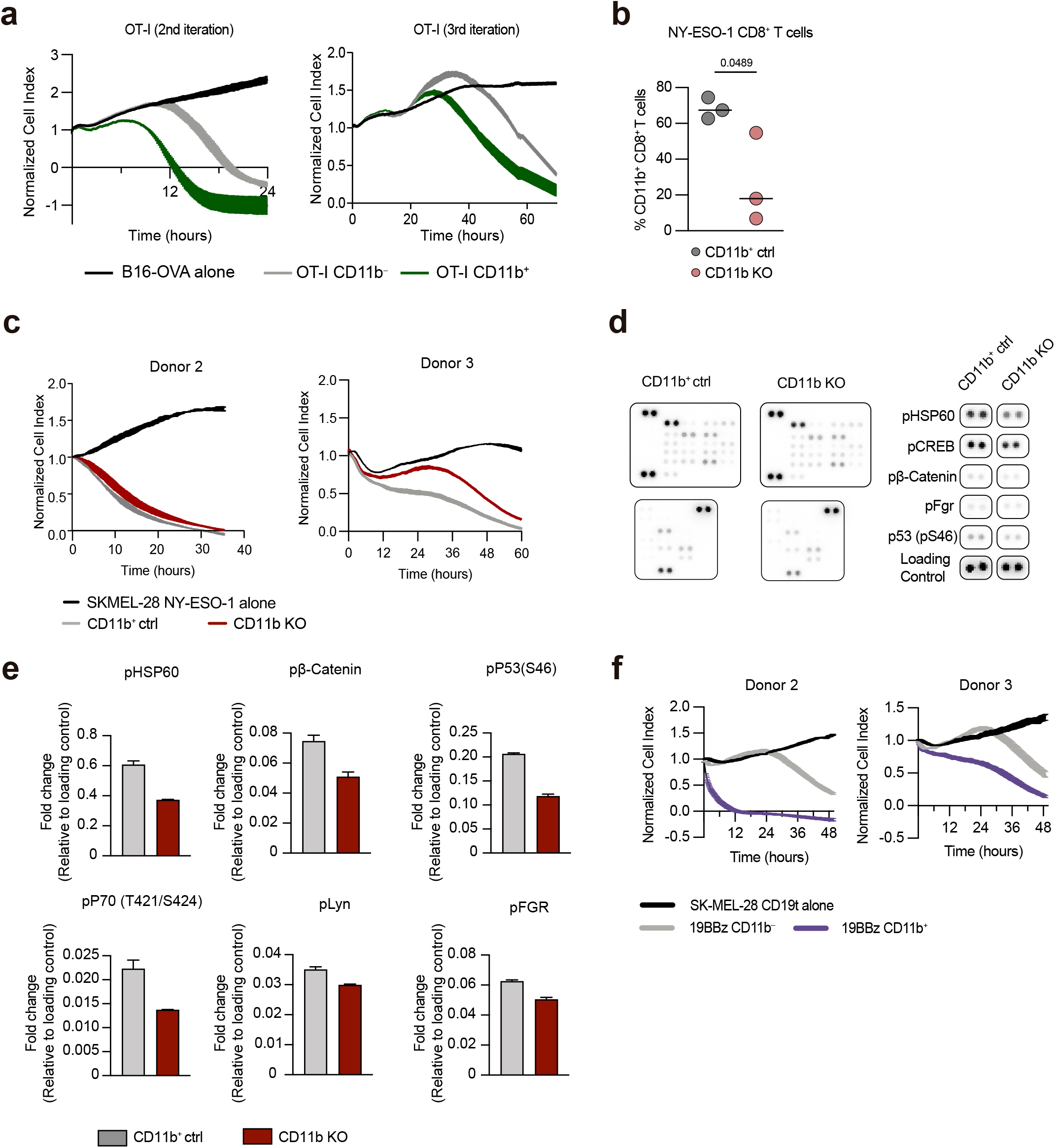
CD11b expression in antigen-specific CD8⁺ T cell cytotoxicity. **a,** Real-time cytotoxicity assay (xCELLigence) evaluating B16-OVA tumor cell lysis when treated OT-I CD11b⁻ or OT-I CD11b⁺ CD8^+^ T cells (E:T = 0.25:1) over a 24 h period. Two additional independent experiments corresponding to Fig. 5b are shown. **b,** Frequency of CD11b⁺ cells among CD11b^+^ control (CD11b^+^ ctrl) and CD11b KO NY-ESO-1-specific human T cells after 10-day in vitro expansion (n = 3 donors). **c,** Analysis of real-time cytotoxicity assay (xCELLigence) evaluating SK-MEL-28 NY-ESO-1 tumor cell lysis when cultured alone, or with control-electroporated (CD11b^+^ ctrl) or CD11b-deficient (CD11b KO) NY-ESO-1 CD8^+^ T cells (E:T = 0.25:1). Two additional independent experiments corresponding to Fig. 5d are shown. **d,e,** Phospho-protein array comparing control-electroporated (CD11b^+^ ctrl) and CD11b-deficient (CD11b KO) human CD8⁺ T cells 4 h after TCR activation. **d**, Full membrane images (left) and selected analytes (right). **e**, Fold changes in signal for the indicated proteins in CD11b KO relative to CD11b^+^ ctrl cells. Duplicate technical spots were averaged before fold-change calculation. One experiment is shown. **f,** Real-time cytotoxicity assay (xCELLigence) evaluating SK-MEL28-CD19t tumor cell lysis when cultured alone or with CD11b^+^ or CD11b⁻ 19BBz CAR-T cells (E:T = 0.25:1). Two additional independent experiments corresponding to Fig. 5f are shown. Data are mean ± s.e.m. (**b**). P values were calculated using a two-tailed, paired Student’s t test (**b**).

